# Unspecific Molecular Adsorption (UMA) sample preparation method for bottom-up and whole protein analysis. The foundation

**DOI:** 10.64898/2026.03.02.709073

**Authors:** Alexandre Zougman

## Abstract

The protein sample preparation methods for shotgun proteomics are nowadays well-established unlike the ones for whole protein analysis. The goal of my work has been to create a simple methodology which provides a single uncomplicated sample preparation tool for these two fields. Nowadays the bulk of proteomics work is done using detergents for protein solubilization. The presented concept, which is based on unspecific adsorption of protein molecules on wide pore materials, allows for protein capture and clean-up from solutions of the most commonly used sodium dodecyl sulfate detergent. It could also be applied to proteins in detergent-free solutions. After the capture and clean-up, proteins could be either cleaved for the downstream peptide analysis or eluted for the whole protein analysis. If required, the eluted whole proteins could be recaptured and cleaved into peptides. Depending on the experimental goals, the sample preparation device could be fitted with embedded proteolytic enzymes to simplify routine sample processing and/or reversed phase media for the downstream peptide or protein separation.

---

Shotgun (bottom-up) protein profiling involves protein cleavage and characterization of the generated peptide products by mass spectrometry (MS) [1]. Also, whole proteins could be profiled directly by MS [2]. Detergents (surfactants) aid in solubilization of proteins as some proteins are either intrinsically hydrophobic or become insoluble in water when denatured. Thus detergents, by solubilizing most of the proteins in biological samples, help provide unbiased coverage for complex protein mixtures. Sodium dodecyl sulfate (SDS), an anionic surfactant, has been the most widely used detergent for protein solubilization for more than 60 years[3] [4]. Several notable methods have been reported over the last years to aid in processing of the SDS-extracted proteins into peptides for the bottom-up proteomics analysis. The Filter Aided Sample Preparation (FASP) method, published in 2009 [5] and of which I have the honor to be a co-author, utilizes membrane ultrafiltration where the membrane pore sizes usually do not exceed 0.01 µm, on-membrane exchange of SDS for urea followed by exchange of urea for a digestion buffer, and on-membrane enzymatic cleavage of proteins into peptides. The Suspension Trapping (STrap) method I co-authored in 2014 [6] involves creation of a fine protein precipitate by acidification of a protein-SDS solution and addition of methanol. A depth filter unit with sub-micron-sized pores (down to ∼ 0.1 µm in size) is consequently used to trap the precipitating protein particles while SDS, which is methanol-soluble, is passing through the filter. After an enzyme addition, the trapped proteins are cleaved into peptides. The Single-Pot, Solid Phase-enhanced Sample-Preparation (SP3) method, also published in 2014, is based on protein capture on functionalized paramagnetic beads in the presence of an organic solvent [7]. After a protein-SDS sample is mixed with a polar organic solvent, during the incubation process the proteins are captured on the beads’ carboxylated surface. The SDS is washed out, an enzyme is added and the proteins are digested into peptides. The mechanism underlying the SP3 protein attachment to the paramagnetic beads was originally thought to be hydrophilic interaction [8] but the phenomenon is likely more complex [9]. While there have been thousands of publications utilizing the mentioned sample preparation tools for the bottom-up protein characterization, reports on streamlined sample preparation for the whole protein MS analysis are scarce. There have been attempts to adjust the established bottom-up sample preparation technologies for use with the whole protein profiling. Dagley et al. performed protein enrichment from HeLa SDS cell lysate using the SP3 technology, eluted captured proteins from the beads with cold 80% formic acid (FA), diluted the eluate 10 times, and analyzed the proteins by reversed phase (RP) chromatography-mass spectrometry [10]. I think it is a fair approach as concentrated formic acid, similarly to SDS, allows for solubilization of most proteins [11], even though the method requires significant dilution of the protein eluate and the use of formic acid could result in undesirable protein modifications [12]. Yang et al. compared membrane ultrafiltration, chloroform-methanol precipitation which was used extensively previously for protein recovery from fractionation experiments [13, 14], and SP3 methodologies for protein clean-up prior to capillary zone electrophoresis-mass spectrometry analysis [15]. E. coli and HepG2 SDS-protein lysates were processed. The captured proteins were recovered with 100 mM ammonium bicarbonate.

The membrane ultrafiltration was found to be superior to the other two tested methods. In my opinion, the ammonium bicarbonate solution alone is not sufficient to completely solubilize a complex proteome which is the definite drawback of the reported approach.

I thought of a new all-inclusive proteomics protein sample preparation tool as a simple device that would capture and clean-up proteins providing the option to straightforwardly process the captured proteins either by cleavage or direct elution. I imagined a solid phase extraction (SPE)-type unit with a wide microporous structure of about 30-μm pore size which would allow for both the quick protein entrapment from SDS-containing solutions and SDS removal as well as the unhindered, low-back-pressure flow-through of the passing fluids facilitating, when required, the fast and efficient downstream whole protein extraction. To optimize the conditions for the protein capture on different porous materials, by trial-and-error visual screening, I used dye-labeled complex protein mixtures solubilized in SDS. Protein labeling with primary-amine-reactive Remazol dyes for gel electrophoresis was demonstrated previously [16, 17]. Even though covalent addition of a dye to a protein may change its properties, such as hydrophobicity, it helps, in my opinion, to visualize general trends in protein mixture behavior during the sample preparation process. I labelled SDS-solubilized protein mixtures with the water-soluble Remazol Brilliant Blue dye. The dye to protein labeling ratio was adjusted to ∼1:8 by weight to allow for partial protein labeling and retaining of some unmodified primary-amine-carrying lysine residues accessible for proteases. As an alternative to the dyed complex protein mixtures, naturally colored proteins could also be used for the sample preparation method testing, such as, for example, cytochrome c with its yellow-red color due to the presence of the heme group.

This approach may be less effective, though, as it does not reflect the protein diversity of a proteome. At the beginning, the screening with the dyed protein lysates had been done using generic tissue paper materials which was followed by testing different types of commercially available wide-pore filter papers. It was found that the addition of a 45% Acetonitrile, 45% Methanol, 100 mM Ammonium Acetate (AMA) solution at the approximate ratio of 6:1 to the protein-SDS sample allowed for complete protein capture by passing the mixture through the tested materials. Incipiently it was noted that the entrapped proteins could be eluted from the filter paper by a concentrated trifluoroethanol (TFE) solution at basic pH (Figure 1). It was also observed that proteins solubilized in this basic TFE solution could be recaptured on a filter paper by addition of 2.5 volumes of acetonitrile and passing the mixture through. The captured proteins could be then digested by trypsin suggesting a way of enzymatic processing for TFE-extracted proteins (Supplementary Figure 1). Even though considered a hazardous substance, TFE has been used in the studies involving protein and peptide solubilization and chromatography for at least 65 years [18–22]. The observation of protein capture on filter papers was confirmed by using protein gel electrophoresis where it was shown that different commercial filter papers with defined pore sizes ranging from 2.7 to 35 μm efficiently captured the loaded protein lysate. The captured proteins were straightforwardly eluted from wide-pore paper filters with basic or acidic solutions of concentrated trifluoroethanol as well as with 8M Urea. The proteins eluted with the acidic TFE solution were successfully recaptured and tryptically digested on strong-cation-exchange (SCX) beads (Figure 2). Also it was found that the TFE-eluted proteins could be captured, under the acidic conditions, by loading onto pre-acidified SG81 Whatman ion exchange paper (which combines cellulose and large pore silica gel) or quartz silica (Supplementary Figure 2) where an enzymatic digest could be performed (Supplementary Figure 3). Urea has been largely used as a protein solubilization agent in two-dimensional gel electrophoresis (2DE) [23]. It is recommended that proteins are reduced and alkylated prior to the first isoelectric focusing step of 2DE [24]. Thus, extraction of proteins with a basic SDS buffer, their denaturation, reduction and alkylation in this buffer, followed by the AMA-aided capture, clean-up and consequent elution with 8M Urea could be a viable simple approach for protein sample clean-up and preparation for 2DE. The protein entrapment under the proposed AMA-loading conditions was not limited solely to the filter paper, other porous materials such as cotton wool, polyester felt, glass wool, nylon, stainless steel mesh and nickel foam were also shown to entrap proteins from SDS lysates (Figure 3). As the protein capture was independent of the chemical composition of the tested capturing materials, the name Unspecific Molecular Adsorption (UMA) for this approach seemed appropriate. Originally, three types of the UMA processing units were suggested – the primary UMA units (tips and mini-columns) incorporating either wide-pore filter paper material or cotton wool and capturing proteins by direct load, the UMA units incorporating porous mesh plugs (e.g. steel mesh) in the tip format suitable for protein capture by pipetting the protein solution up and down through the plugs several times, and the UMA reversed phase units composed of the upper wide-pore filter paper part and bottom reversed phase part suitable to directly capture and, consequently, release the peptides/proteins eluted from the upper part (Supplementary Figure 4). The loaded and cleaned-up proteins could be quickly digested *in situ* with introduced trypsin at elevated temperature, with the digest products eluted for downstream analysis (Figures 4, 5 and Supplementary Figure 5). Instead of digestion, the captured proteins could be eluted with acidic solutions containing trifluoroethanol (Figure 6). These eluted proteins could be analyzed directly by positive electrospray-mass spectrometry as demonstrated by MS analysis of SDS-solubilized standard proteins processed using different UMA units with the elution by either 35% or 70% TFE in 1% FA. The eluted proteins were loaded into emitter needles and analyzed in a positive nanospray mode on a Q-TOF mass spectrometer (Figure 7, Supplementary Figures 6 and 7). In addition, the eluted proteins could be chromatographically profiled by loading onto wide pore reversed phase columns suitable for protein separations as shown in Figure 8. The figure shows an LC-MS profile of an SDS-solubilized protein standard mixture, with protein masses ranging from ∼6 to ∼66 kDa (insulin, trypsin, carbonic anhydrase and albumin), which was processed on an UMA paper tip. The captured proteins were eluted with 35% TFE in 1% FA. The undiluted protein eluate was loaded onto and separated on a C4, 300 Å capillary column using increasing gradient of acetonitrile/isopropanol mixture in 0.1% trifluoroacetic acid with protein MS profiling done by an orbitrap mass spectrometer. It was observed as well, while working with simple protein mixtures of water-soluble proteins, such as antibodies, that elution with a diluted acid solution (e.g. 0.1% TFA) without trifluoroethanol was sufficient for extraction of the captured proteins from UMA paper tips and their consequent MS analysis. A monoclonal antibody solution was either used as is or had SDS added to it to the final concentration of 4%. The antibody samples were processed on UMA paper tips and eluted with 0.1% TFA. The antibody eluates were successfully profiled by LC-MS using a time-of-flight MS system (Suppplementary Figure 8).

**Figure 1.**
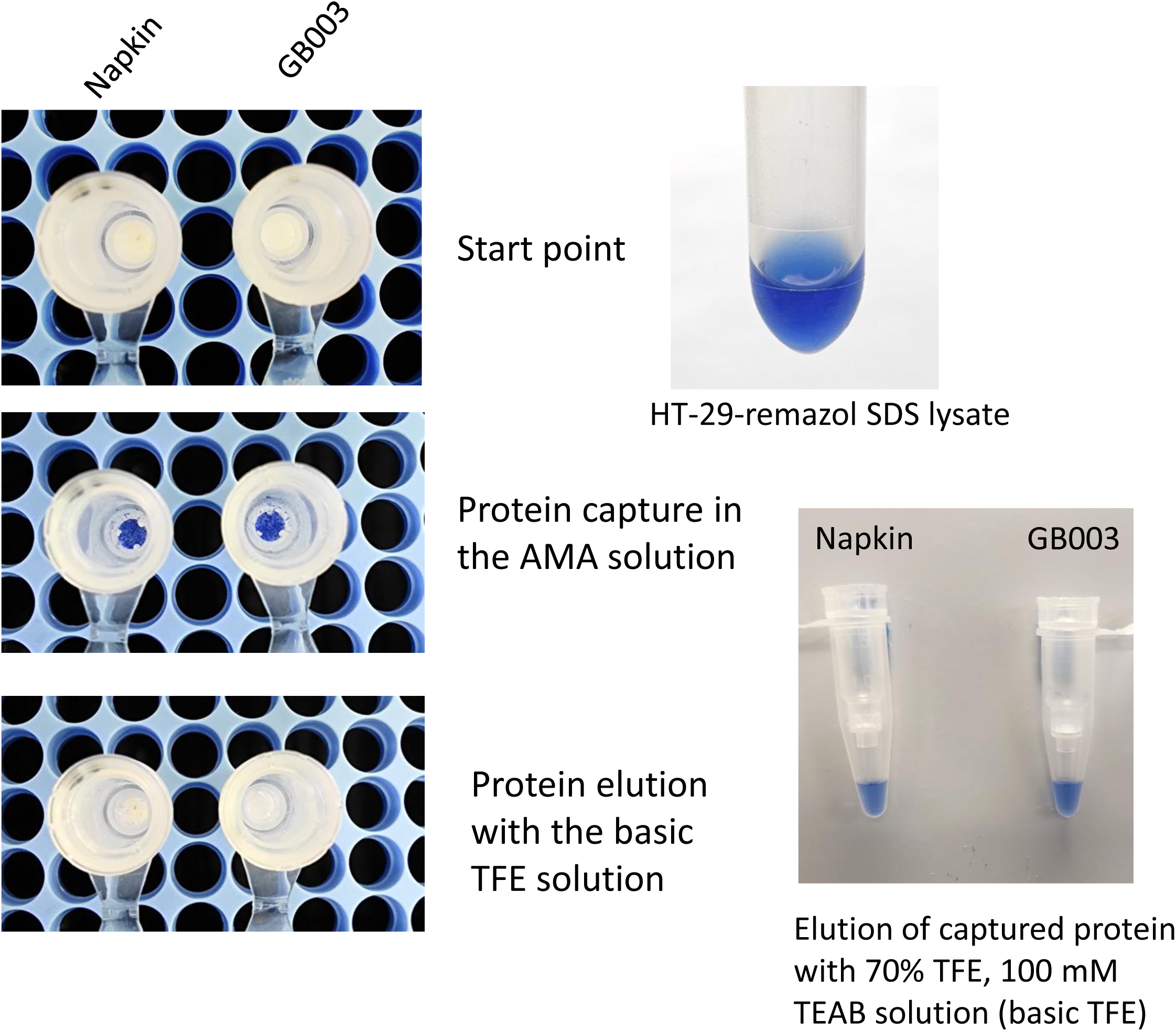
Dyed protein capture and elution from UMA paper mini columns. UMA mini units were prepared either with a generic paper napkin or GB003 (Whatman) filter paper. HT-29-remazol-dyed protein SDS lysate was loaded into the units in 6 volumes of the AMA solution (45% acetonitrile, 45% methanol, 100 mM ammonium acetate). Following the AMA wash, the captured proteins were eluted with the basic TFE solution (70% TFE, 100 mM TEAB).

**Figure 2.**
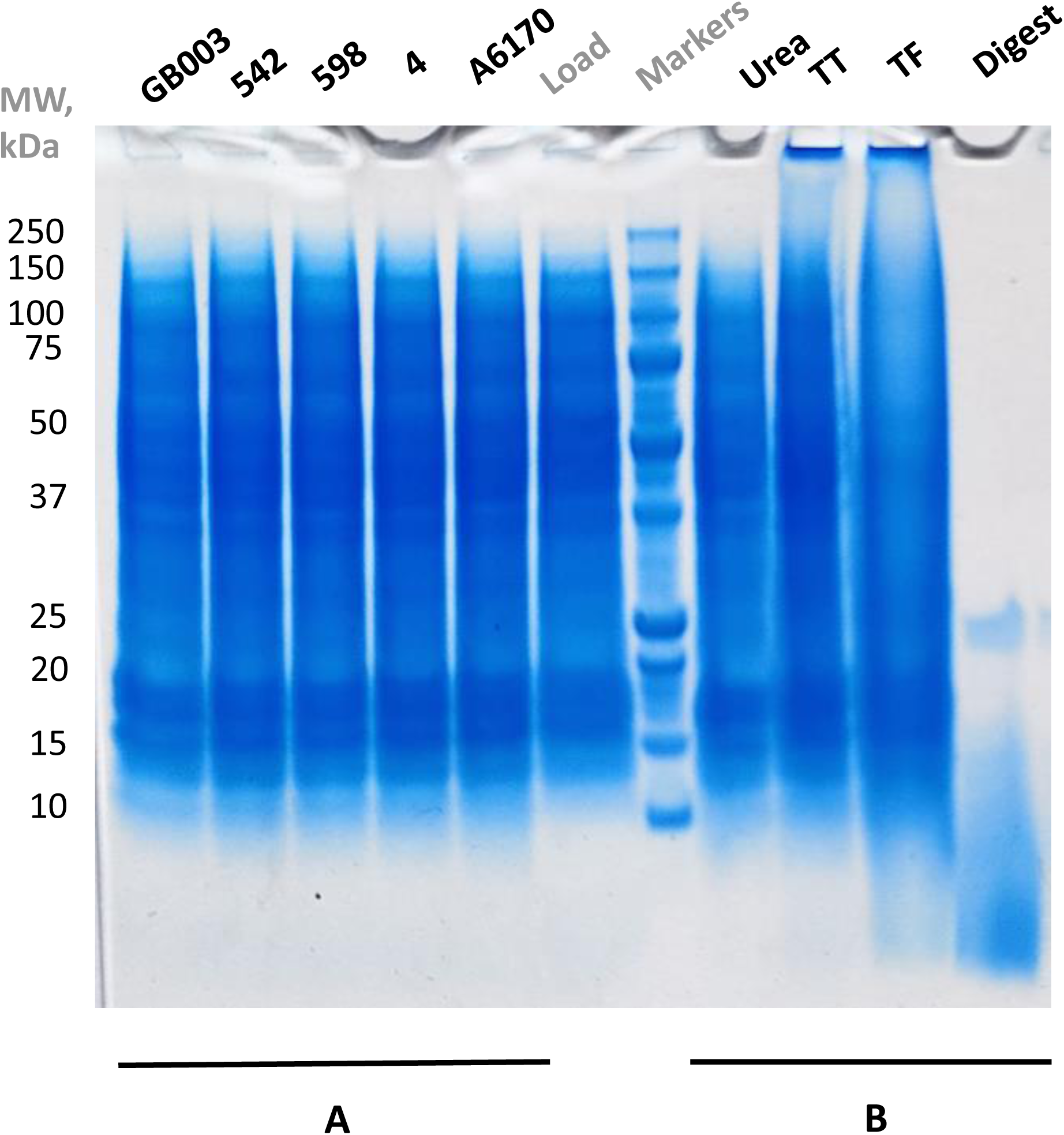
Testing protein capture and extraction on different commercial filter papers using UMA mini units and visualization by gel electrophoresis. Testing acidic-TFE-extracted-protein digestion on SCX beads. **A.** ∼50 μg of HT-29 cell lysate in 4% SDS, 50 mM Tris-HCl, pH 7.6, was loaded in 6 volumes of the AMA solution into the UMA mini units made with different filter papers - Ahlstrom 6170 (35 μm pore), Whatman 4 (20-25 μm pore), Whatman 598 (8-10 μm pore), Whatman 542 (2.7 μm pore), Whatman GB003 (wide pore, unspecified pore size). The captured proteins were eluted with 1X Laemmli buffer. **B.** ∼50 μg of HT-29 cell lysate in 4% SDS, 50 mM Tris-HCl, pH 7.6, was loaded into the GB003 paper UMA mini units. The captured proteins were eluted either with 8M Urea, 70% TFE in 100 mM TEAB (TT) or 70% TFE in 1% Formic Acid (TF). In addition, the TF-eluted proteins were recaptured on an SCX tip, washed with water and digested with trypsin in 100 mM AmBic. The urea-eluted proteins were concentrated, washed on a 3 kDa centrifugal filter and solubilized in Laemmli buffer prior to gel electrophoresis. The TT- and TF-eluted proteins as well as the digest products were dried down and resolubilized in 1X Laemmli buffer. The samples were run on a 4-12% Bis-Tris protein gel. The gel was stained with Coomassie.

**Figure 3.**
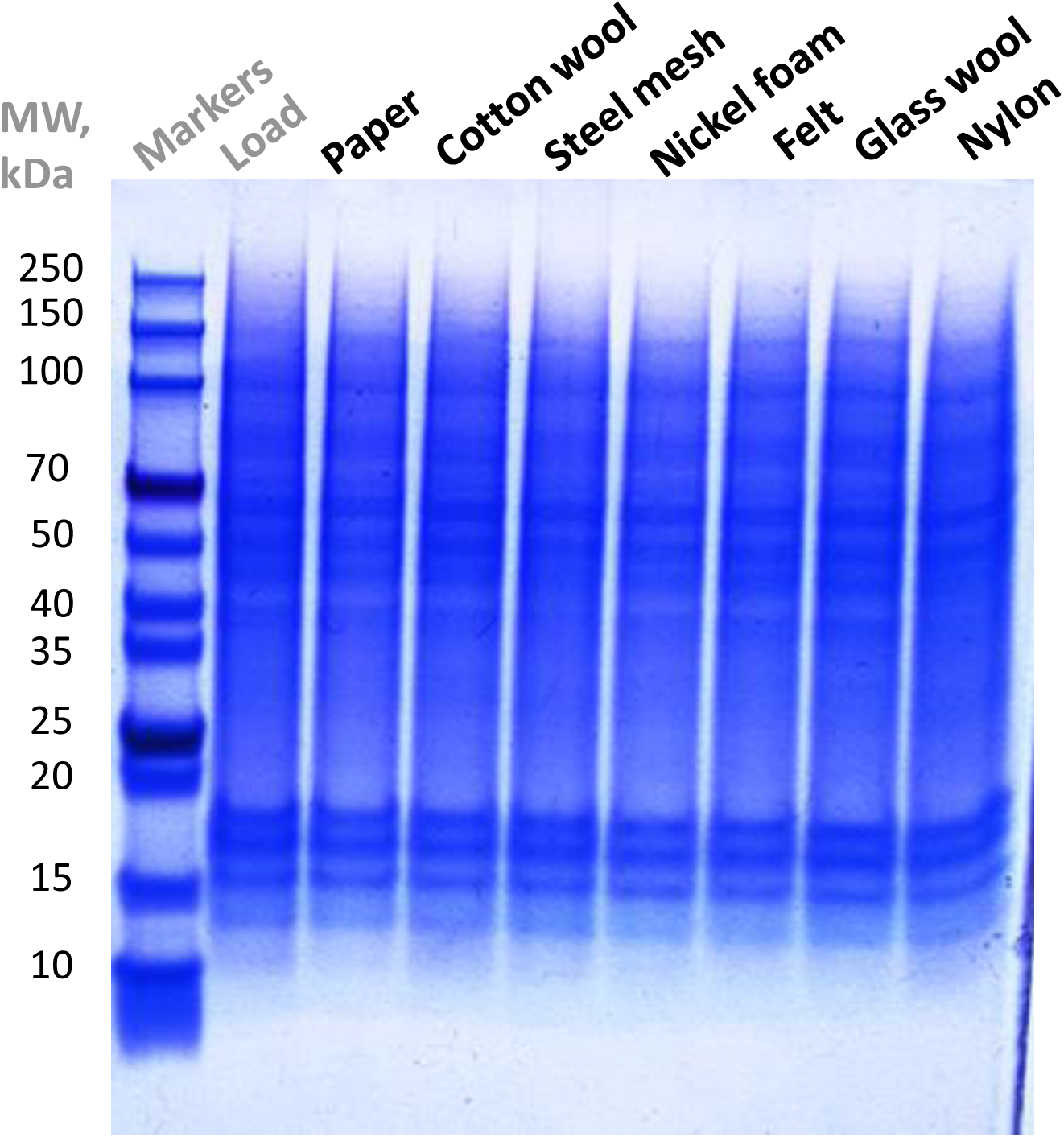
Testing the AMA-assisted protein capture on different materials. Visualization by gel electrophoresis. HT-29 SDS cell lysate proteins were captured by UMA tips with different plugs utilizing the AMA loading approach (1 sample volume:6 AMA volumes). Paper, cotton wool, polyester felt, glass wool, nylon tips were loaded directly by passing the AMA-lysate mix through the tips; steel mesh and nickel foam were loaded by pipetting the AMA-lysate mixture up and down the corresponding plugs 30 times. After the AMA wash, the captured proteins were eluted with 1X Laemmli buffer. The samples were run on a 4-12% Bis-Tris protein gel. The gel was stained with Coomassie.

**Figure 4.**
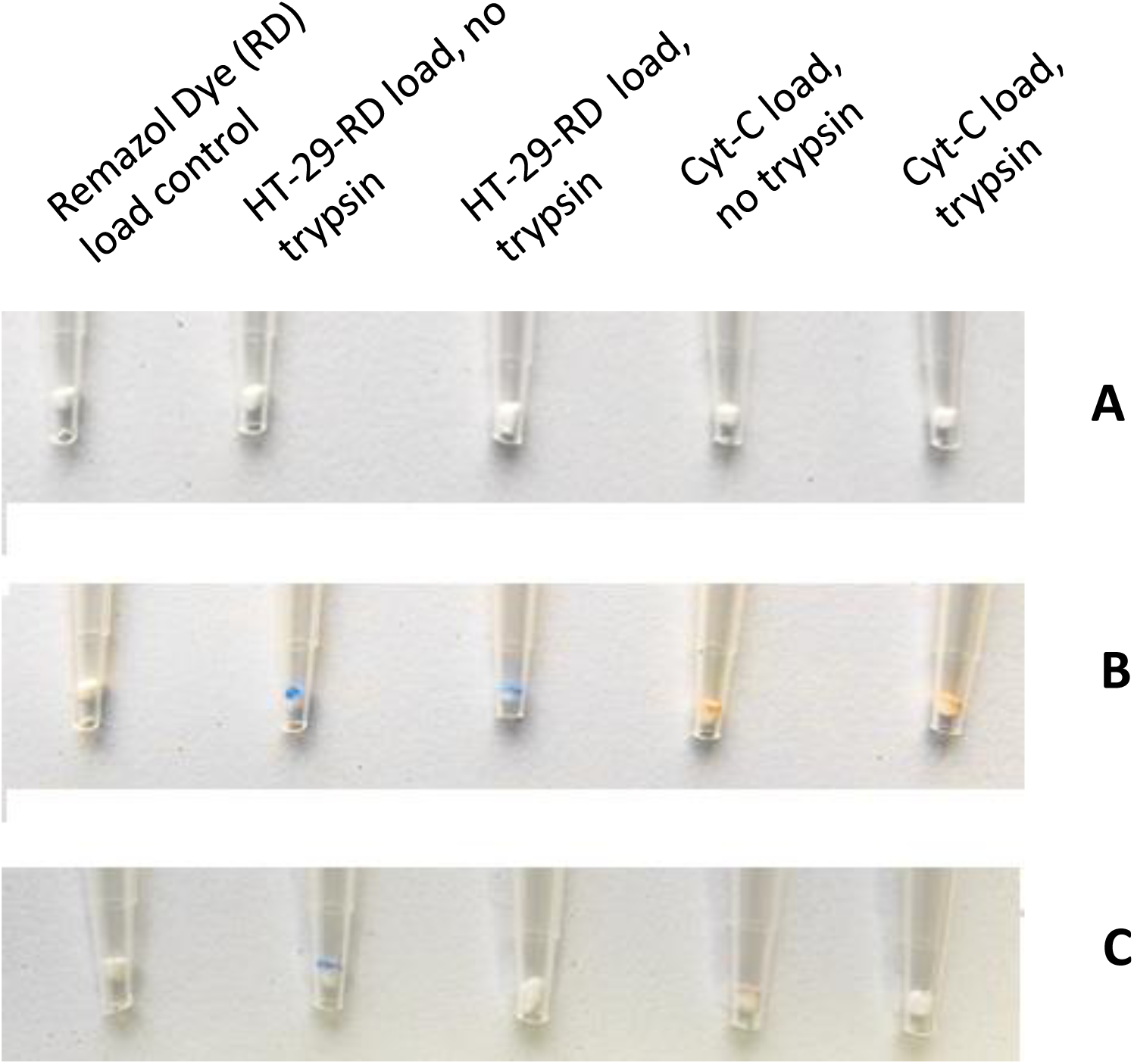
Protein capture and digestion on UMA paper tips using dyed Remazol HT-29 SDS protein lysate and Cytochrome-C SDS solution. UMA tips were constructed utilizing GB003 filter paper **(A)**. The UMA tips were loaded with either Remazol Dye (RD), HT-29-RD dyed cell lysate or Cytochrome-C in 4% SDS in 6 volumes of the AMA solution. To the captured HT-29-RD or Cytochrome-C proteins either trypsin in 70 mM AmBic or only AmBic was added followed by incubation at 56 °C for 30 min **(B)**. The samples were eluted after incubation **(C)**.

**Figure 5.**
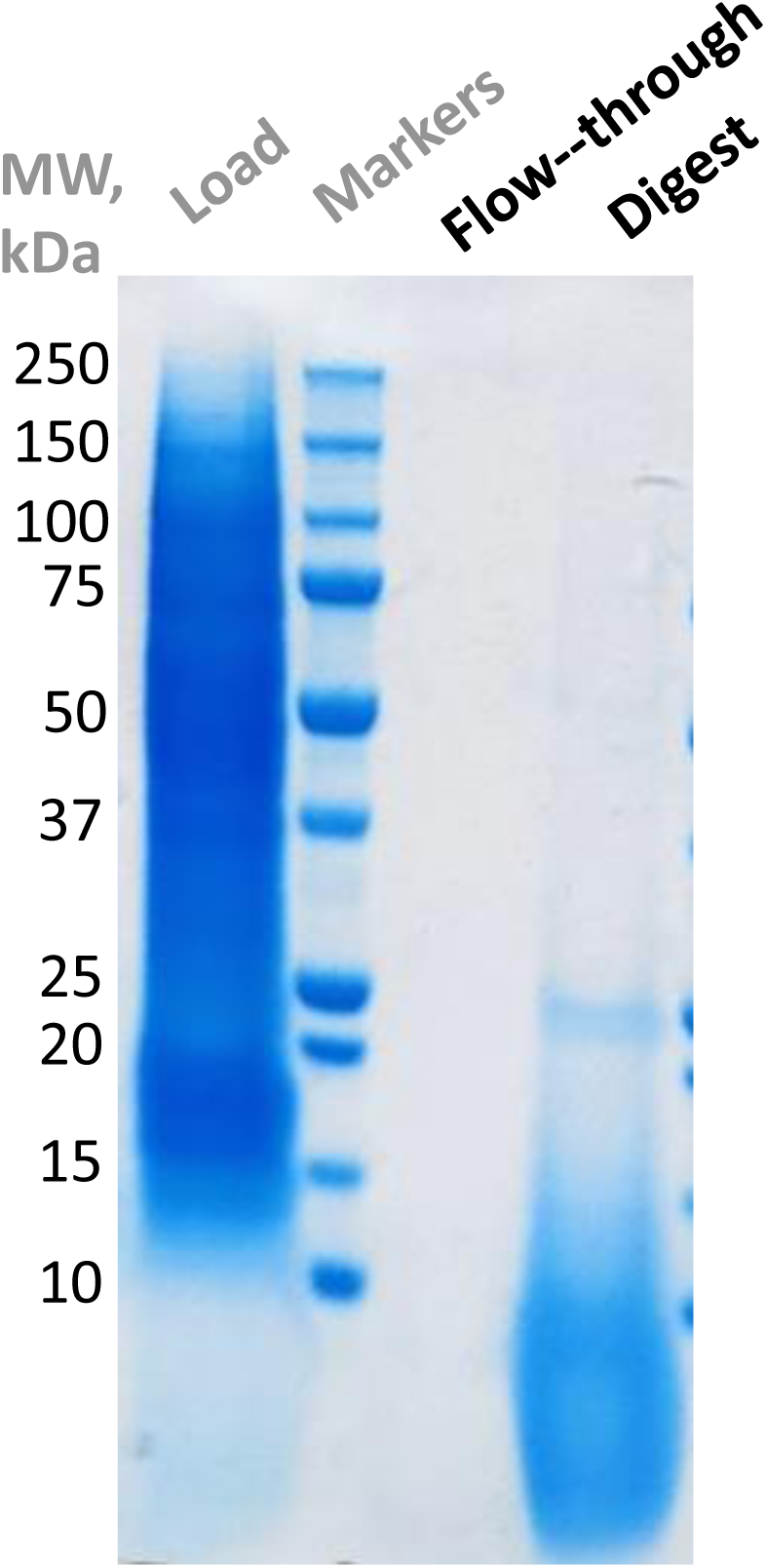
Protein capture and digestion on an UMA paper mini column visualized by gel electrophoresis. ∼50 μg of HT-29 cell protein lysate in 4% SDS, 50 mM Tris-HCl, pH 7.6, was processed using an UMA mini unit. The lysate proteins were reduced, alkylated and loaded into the UMA unit in 6 volumes of the AMA solution. After the wash with the AMA solution, tryptic digest was performed for 45 min at 53 °C in 70 mM AmBic. The digest products were eluted with 1X Laemmli buffer. The samples were run on a 4-12% Bis-Tris protein gel. The gel was stained with Coomassie.

**Figure 6.**
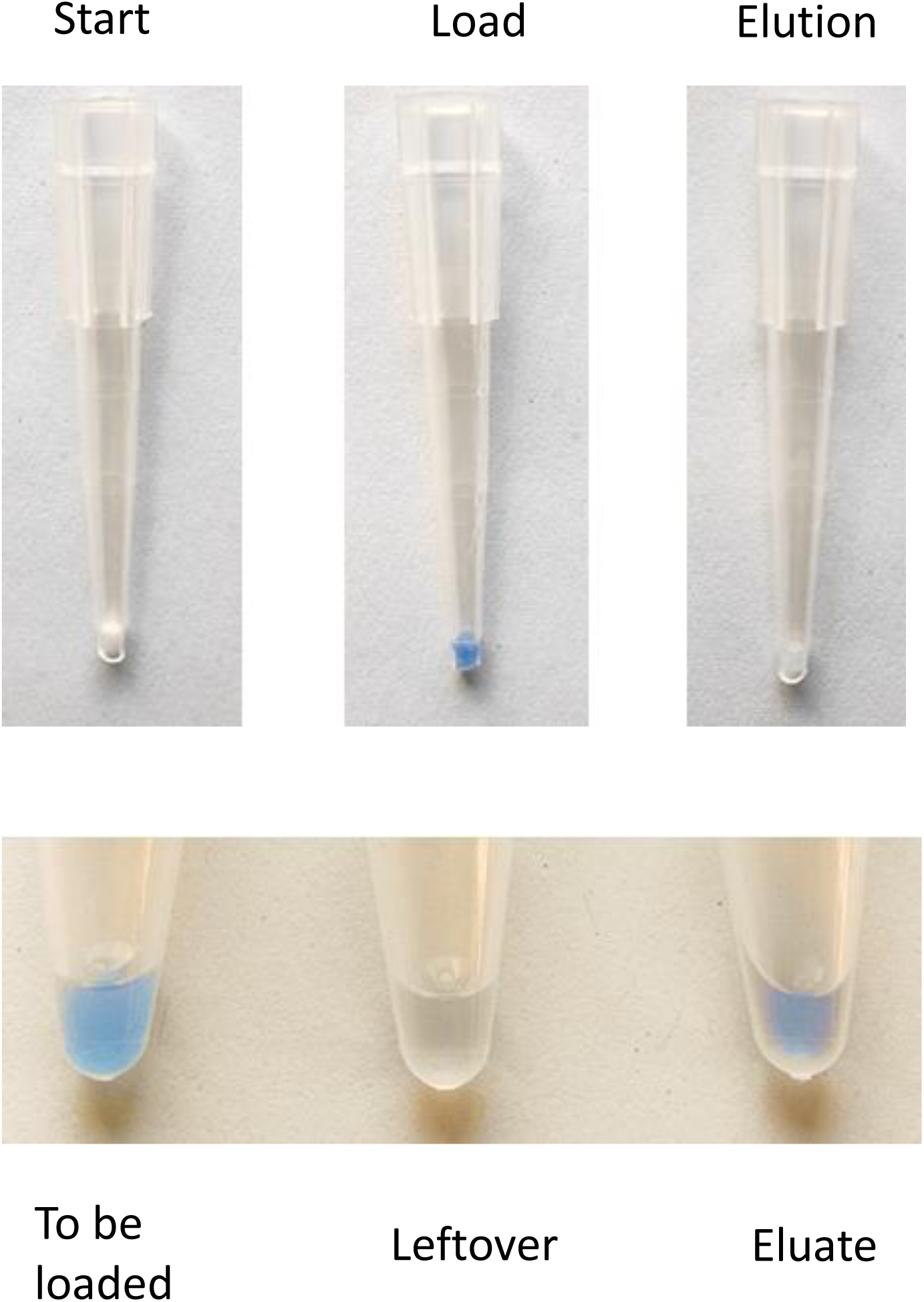
Capture and elution of dyed serum proteins using an UMA pipetting tip. ∼20 μg of Remazol-dyed serum proteins in 4% SDS, 50 mM NaHCO3 buffer were mixed with five volumes of the AMA solution. The mix was loaded into a loosely packed cotton wool tip by pipetting up and down several times. The bound proteins were washed with the AMA solution and eluted with 70% TFE in 1% FA.

**Figure 7.**
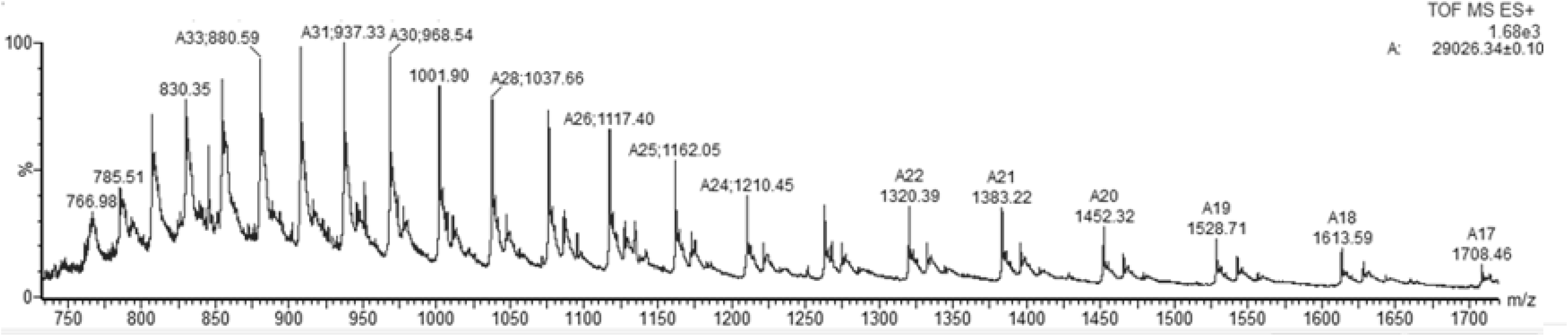
MS analysis of Carbonic Anhydrase solubilized in SDS, cleaned up and eluted using an UMA tip with a cotton wool plug. Carbonic Anhydrase (C3934, Sigma). Predicted MW 29,024.6 Da with N-term Met removed and N-term acetylated. The observed delta mass of +2 Da could correspond to an amino acid substitution (e.g. Leu/Ile->Asp). The protein was dissolved in 4% SDS, 50 mM TEAB buffer. An UMA tip with a cotton wool plug was used. Six volumes of the AMA solution were added to the protein solution aliquot containing 10 μg of protein. The mixture was loaded into the UMA tip by centrifugation. After a wash with the AMA solution, the protein was eluted in 35% TFE, 1% FA. The eluate was analyzed by nanospray in the positive ionization mode on Synapt G1 Q-TOF mass spectrometer.

**Figure 8.**
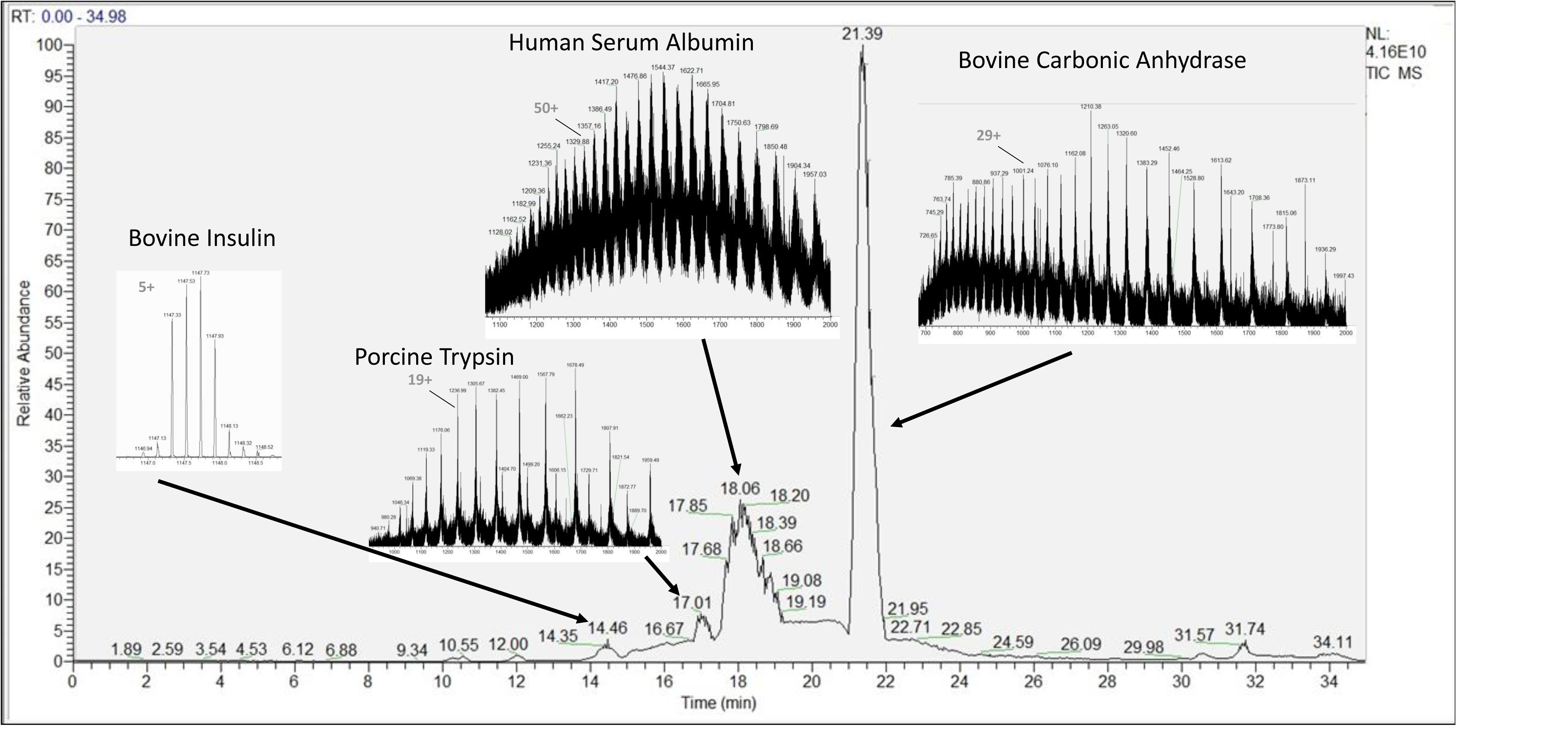
LC-MS elution profile of a 4-protein SDS mixture captured, cleaned-up and eluted using an UMA paper tip. Bovine insulin (I6634, Sigma), porcine trypsin (T0303, Sigma), bovine carbonic anhydrase (C3934, Sigma) and human serum albumin (A3782, Sigma) were solubilized in 4% SDS, captured and cleaned up in an UMA paper tip with the AMA solution, and then eluted with 35% TFE in 1% FA. The proteins were loaded undiluted onto and separated on a C4 capillary column. The MS spectra were acquired in the orbitrap part of LTQ-Orbitrap Velos mass spectrometer.

I compared the UMA approach with the established STrap methodology for protein clean-up and digestion [6]. Since its publication and commercialization, the STrap concept has been widely used in thousands of publications covering the broad area of the bottom-up-based proteomics research, with samples as diverse as plants, bacteria, mammalian cells and tissues, body fluids and archaeological material. Performance of the UMA method using direct load (UMA tips) and pipetting (UMA-P tips) units was compared to the STrap method (STrap tips with quartz filter plugs, STrap processing protocol [6]) for the analysis of a PC3 cell protein lysate by tryptic digestion. The preliminary visualization by gel electrophoresis showed similar digest outputs for all tested approaches (Figure 9). The protein lysate was then processed by the UMA, UMA-P and STrap units using trypsin digestion for 45 min at 53 °C in four technical replicates. The resultant peptides were analyzed by LC-MS/MS on an orbitrap mass spectrometer. The data analysis results showed a slightly better average performance of the UMA and UMA-P approaches compared to the STrap in terms of total number of proteins identified. But, nevertheless, the methods performances seemed very similar (Figure 10, Supplementary Table 1).

**Figure 9.**
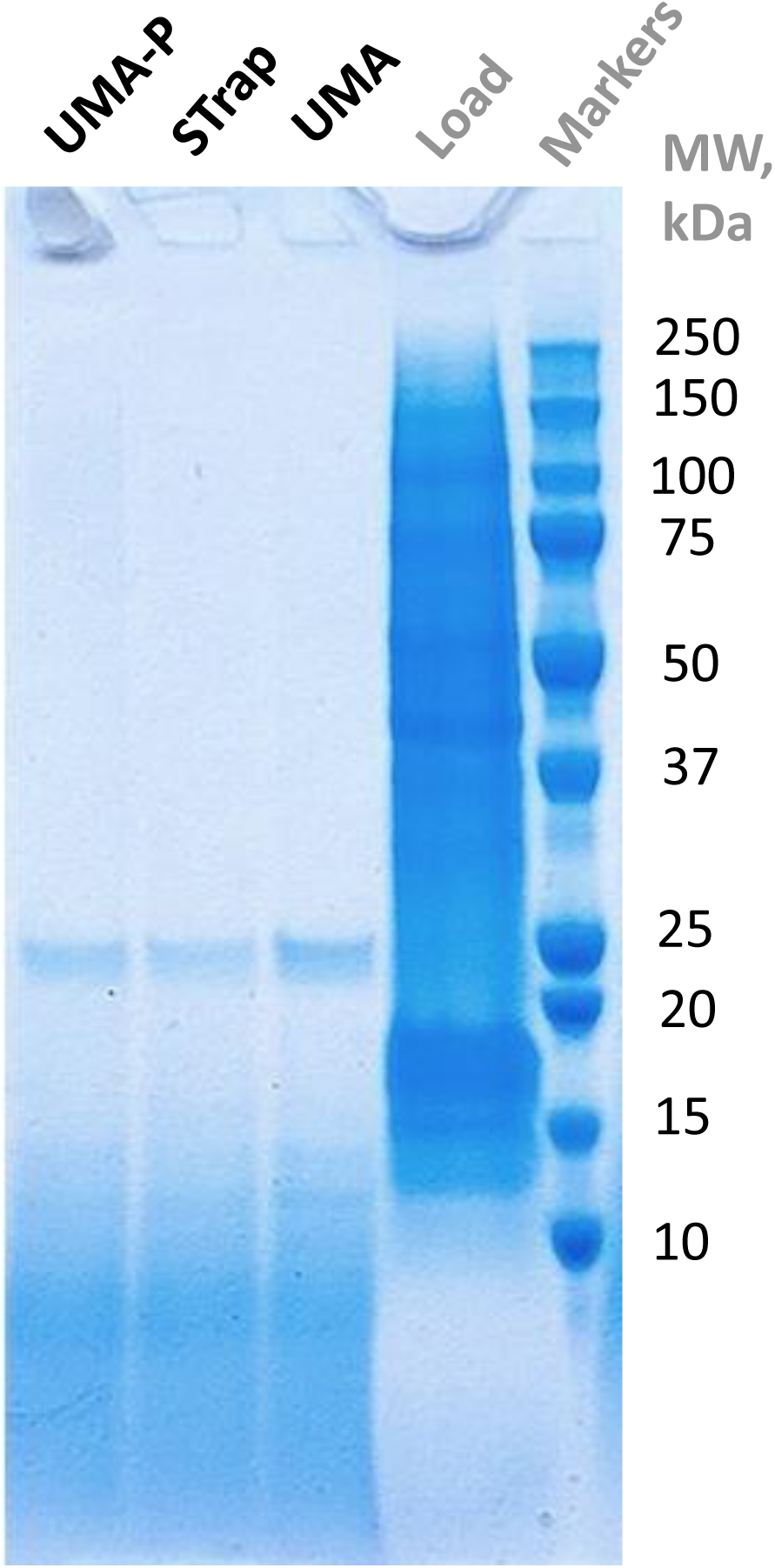
Proteolytic processing of PC3 cell SDS protein lysate with STrap, UMA direct loading (UMA) and pipetting (UMA-P) units. Visualization by gel electrophoresis. ∼50 μg of PC3 cell protein lysate in 4% SDS, 50 mM Tris-HCl, pH 7.6, was processed using STrap, UMA (direct loading, paper) and UMA-P (pipetting, steel mesh) tips with relevant protocols. Tryptic digest was performed for 30 min at 53 °C in 70 mM Ammonium Bicarbonate. The digest products were eluted in 70 mM AmBic. The samples were run on a 4-12% Bis-Tris protein gel. The gel was stained with Coomassie.

**Figure 10.**
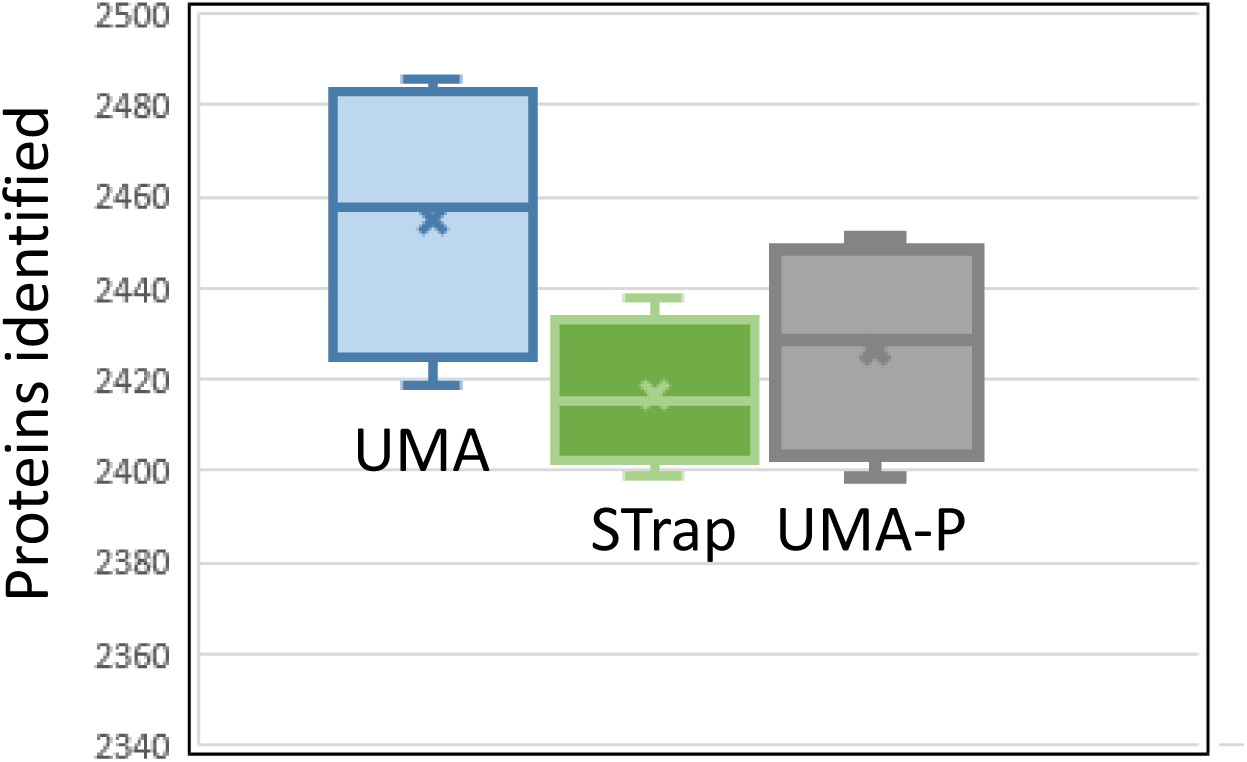
Proteolytic processing of PC3 cell SDS protein lysate with STrap, UMA direct loading (UMA) and pipetting (UMA-P) units. Analysis by LC-MS/MS. PC3 SDS cell protein lysate was processed by the UMA, STrap and UMA-P approaches using trypsin digestion for 45 min at 53 °C in four replicates. The resultant digest products were analyzed twice using capillary chromatography on LC-MS Exploris 240 orbitrap system. The total acquisition time for each run was 45 min. The data were processed by MaxQuant (https://maxquant.net/). For protein identification at least one unique peptide per protein was required.

To check the applicability of the new UMA methodology in relation to the analysis of low-amount less complex protein samples, I tested the viability of the rabbit anti-FOXM1 antibody after an extended long-term storage period (∼10 years at 4°C). I performed an immunoprecipitation experiment using either 2 μg of this antibody or a control rabbit antibody together with 1 mg of HeLa protein extract, in triplicates. Bound proteins were eluted with 4% SDS and processed using the UMA approach, digestion by trypsin and consequent LC-MS/MS analysis. The targeted FOXM1 protein was clearly identifiable in the tested anti-FOXM1 immunoprecipitates (Supplementary Figure 9, Supplementary Table 2).

Comparably to the previously reported [25] I suggested that abundant serum proteins would not bind to the materials used in UMA pipetting units in a basic buffer but some less abundant proteins would bind to these materials. It was later confirmed by gel electrophoresis – the abundant serum proteins did not seem to be binding to the UMA-P steel mesh tips after solubilization in the 250 mM Triethylammonium bicarbonate (TEAB), 5 mM Tris(2-carboxyethyl)phosphine (TCEP), 10 mM Iodoacetamide (IAA) basic binding buffer solution (proteins are reduced and alkylated while in this binding buffer). The addition of several volumes of acetonitrile to this basic serum solution induced the binding of the serum abundant proteins to the UMA-P tips (Supplementary Figure 10). This simple serum fractionation approach was compared to the direct analysis of the unfractionated serum.

The serum was processed either by fractionation on the UMA-P tips (fraction 1 – protein capture from the basic serum solution, fraction 2 – capture after addition of two volumes of acetonitrile, fraction 3 – capture after addition of another volume of acetonitrile) or by direct UMA preparation using trypsin digestion for 45 min at 53°C, in three replicates. The digests were analyzed by LC-MS/MS. As expected, the fractionation approach noticeably increased the number of identified proteins (Supplementary Figure 11, Supplementary Table 3).

It was a random find while working with elastase that this enzyme could be attached to the filter paper or cotton wool plug surface from its diluted solutions in deionized water (the enzyme concentration was ∼ 0.01 μg/μl) either by slowly passing the elastase solution through a filter paper or cotton wool plug in a tip or incubating the elastase solution in deionized water with a filter paper piece. While not the first choice for complex proteome profiling due to its unspecific ladder-like digestion pattern, elastase could actually be the right enzyme for the sequence characterization of a single protein or protein mixture of low complexity. Several standard proteins - bovine insulin, porcine trypsin, bovine carbonic anhydrase and human serum albumin - were solubilized in 4% SDS, reduced and alkylated, and loaded using the AMA approach onto the UMA units containing the embedded elastase. After incubation in 70 mM ammonium bicarbonate (AmBic) at 57 °C for 30 min the resultant peptide products were analyzed by LC-MS/MS. The identified peptide ladders allowed for complete or nearly complete protein sequence coverage maps. I also found that the ladder-like protein cleavage pattern could be obtained by incubating the UMA-captured proteins in a solution of a strong acid at 85-90 °C for about 30 min in the optional presence of TCEP as a reducing agent (e.g. 10% hydrochloric acid, 50 mM TCEP). For this purpose UMA-C_18_ units were made which contained a filter paper on the top, used for protein capture and clean-up, and C_18_ reversed phase media at the bottom, used for the capture, clean-up and elution of the peptide products of the acidic protein cleavage. The optional use of the reducing agent during the acidic chemical cleavage process removes the need for preliminary reduction and alkylation of the introduced proteins. Because the protein cleavage, reduction, clean-up and elution steps are performed in the acidic environment, there is no risk in the disulfide bond reformation. Similarly to the elastase digest, the acidic cleavage of the standard proteins resulted in the ladder-like peptide patterns and complete or nearly complete protein sequence coverage (Figure 11 and Supplementary Figure 12). It was observed that the acidic cleavage resulted in deamidation of asparagine and glutamine residues (and if a protein was alkylated with iodoacetamide – in deamidation of carboxyamidomethyl cysteine, i.e. carboxyamidomethyl conversion to carboxymethyl (Supplementary Figure 13)). I see the described elastase and acidic cleavage approaches as simple and useful tools for complete protein sequence determination.

**Figure 11.**
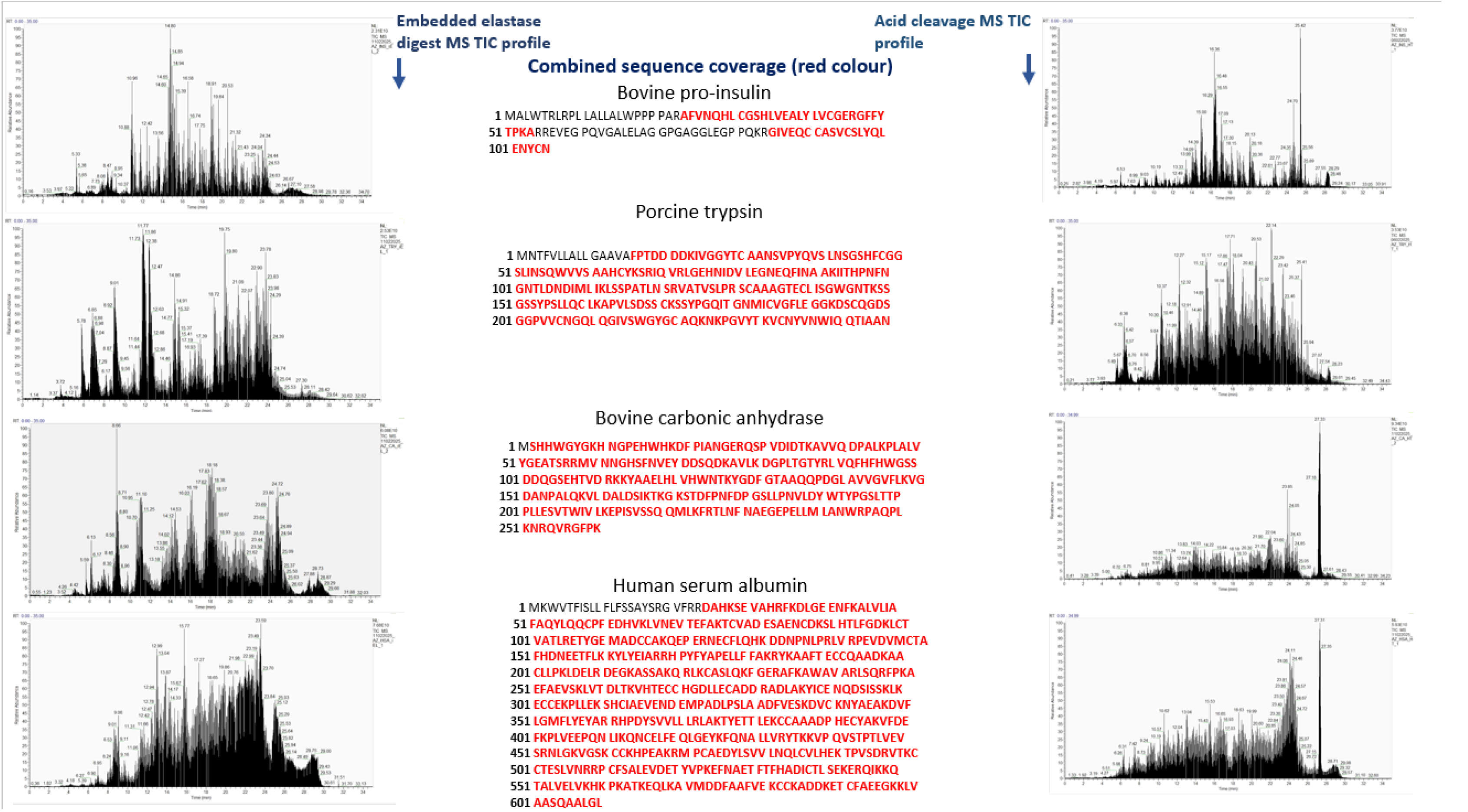
LC-MS/MS sequence coverage of standard proteins either digested in embedded elastase UMA units or cleaved by HCl-TCEP using UMA-C18 units. Bovine insulin (I6634, Sigma), porcine trypsin (T0303, Sigma), bovine carbonic anhydrase (C3934, Sigma) or human serum albumin (A3782, Sigma) were separately solubilized in the 4% SDS, 50 mM TEAB buffer. The proteins were processed using either digestion in embedded-elastase UMA units or acidic cleavage in UMA-C18 units. The resulting peptide products were analyzed by LC-MS/MS on LTQ-Orbitrap Velos mass spectrometer.

Similarly to this elastase observation, it turned out that other enzymes such as trypsin and chymotrypsin could also be adsorbed on a filter paper or cotton wool from their diluted solutions in deionized water. The adsorbed enzymes also seemed to be capable of digesting proteins in the UMA settings immediately after the adsorption (Figures 12 and Supplementary Figure 14). Further tests ensued. An UMA paper tip was ‘charged’ with trypsin by slowly passing the diluted trypsin solution in deionized water through it followed by washes with water and ethanol. The embedded-trypsin UMA tip was left for two weeks at room temperature (RT) prior to the experiment. The HT29-cell SDS protein lysate dyed with the Remazol Blue dye was loaded into the UMA-trypsin tip. After a short incubation at 53 °C in the 70 mM ammonium bicarbonate digestion buffer, it was clear that the captured proteins were cleaved as it became possible to recover the entrapped matter by elution in the digestion buffer (Figure 13). The viability of this approach was again confirmed by gel electrophoresis of the peptide products obtained using a prepared UMA mini column unit with embedded trypsin which had been stored at RT for two weeks prior to the experiment. The HT29-cell SDS lysate was loaded into this unit with the AMA solution. Incubation for 30 min at 53 °C in the AmBic digestion buffer was performed and the digest products were eluted from the unit. The gel picture confirmed full capture of the loaded proteins prior to the digest and the total protein cleavage after the digest completion (Figure 14). A similar picture was observed when digesting alcohol dehydrogenase protein preparation labeled with the Remazol Blue dye (Supplementary Figure 15). The protein digestion in the embedded-trypsin units was fast at the elevated temperature and, visually, there was no difference observed on the gel between the peptide outputs after 10-min, 20-min and 30-min incubations. The embedded-elastase units did not show such a straightforward trend (Figure 15). The performance of the embedded trypsin approach (UMA-T) was compared to the original UMA concept for processing of sds-solubilized cellular lysates. The PC3-cell protein lysate was digested either with the UMA-T units (the units were stored for two weeks at RT prior to the experiment) or the UMA units in four technical replicates for 45 min at 53 °C. The digests were analyzed by LC-MS/MS on an orbitrap mass spectrometer. There was no significant difference observed in the number of identified proteins (Supplementary Figure 16, Supplementary Table 4) which suggested that the UMA units with embedded enzymes could be a practical alternative to the units requiring conventional enzyme introduction. Also, it was observed subsequently that the enzyme adsorption under the described conditions was not limited to the paper or cotton wool materials and could also be performed on other materials such as, for example, silica or polyester (Supplementary Figure 17).

**Figure 12.**
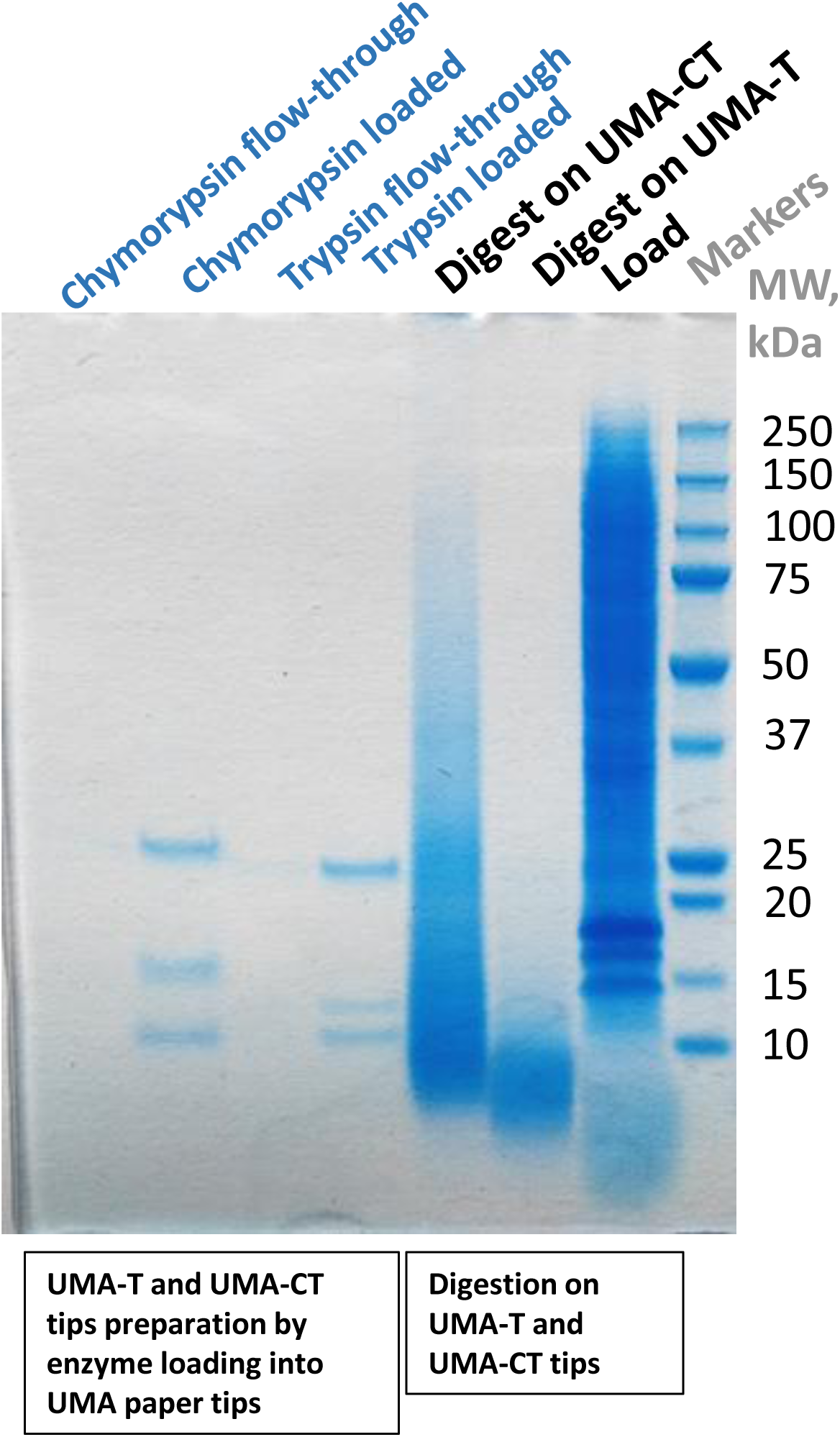
Trypsin and chymotrypsin ‘charging’ of UMA paper tips. Embedded enzyme digestion of 293T cell SDS protein lysate on the ‘enzyme-charged’ tips. Visualization by gel electrophoresis. UMA paper tips were constructed by placing two plugs of the GB003 paper filter into 200-μl pipette tips. The UMA-T tip was prepared by passing ∼2 μg of trypsin in 180 μl of deionized water through the UMA tip and then washing the UMA tip with deionized water and ethanol. The UMA-CT tip was prepared the same way using chymotrypsin. 293T SDS cell protein lysate was loaded into the UMA-T and UMA-CT tips in 6 volumes of the AMA solution. After the wash with the AMA solution, the digestion was performed at 53 °C for 30 min in 70 mM AmBic. The digest products were eluted with 1X Laemmli buffer. The samples were run on a 4-12% Bis-Tris protein gel. The gel was stained with Coomassie.

**Figure 13.**
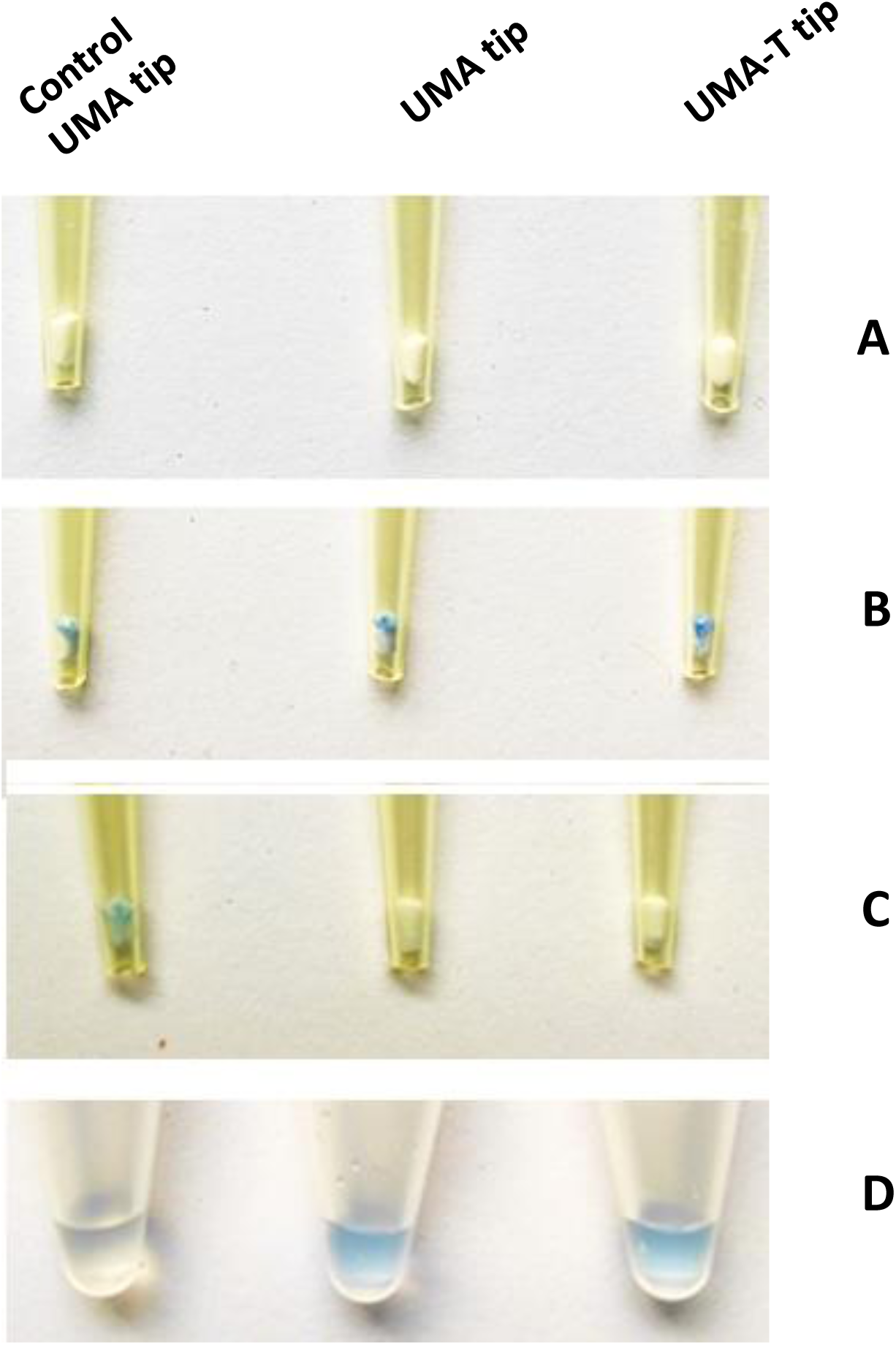
Digestion of dyed HT-29 SDS protein lysate on UMA tips using either post-protein-capture trypsin loading or embedded trypsin. UMA paper tips were constructed by placing plugs of the GB003 filter paper into 200-μl pipette tips. The UMA-T tip was prepared by passing 2.5 μg of trypsin in 200 μl of deionized water through the UMA paper tip and then washing this tip with deionized water and ethanol **(A)**. HT-29-remazol protein lysate was loaded in and washed with the AMA solution **(B)**. Control and UMA-T-tip – incubation with 70 mM AmBic, UMA tip – incubation with trypsin in 70 mM AmBic. Post-incubation elution was done with 70 mM AmBic **(C)**. Post-incubation eluates **(D)**.

**Figure 14.**
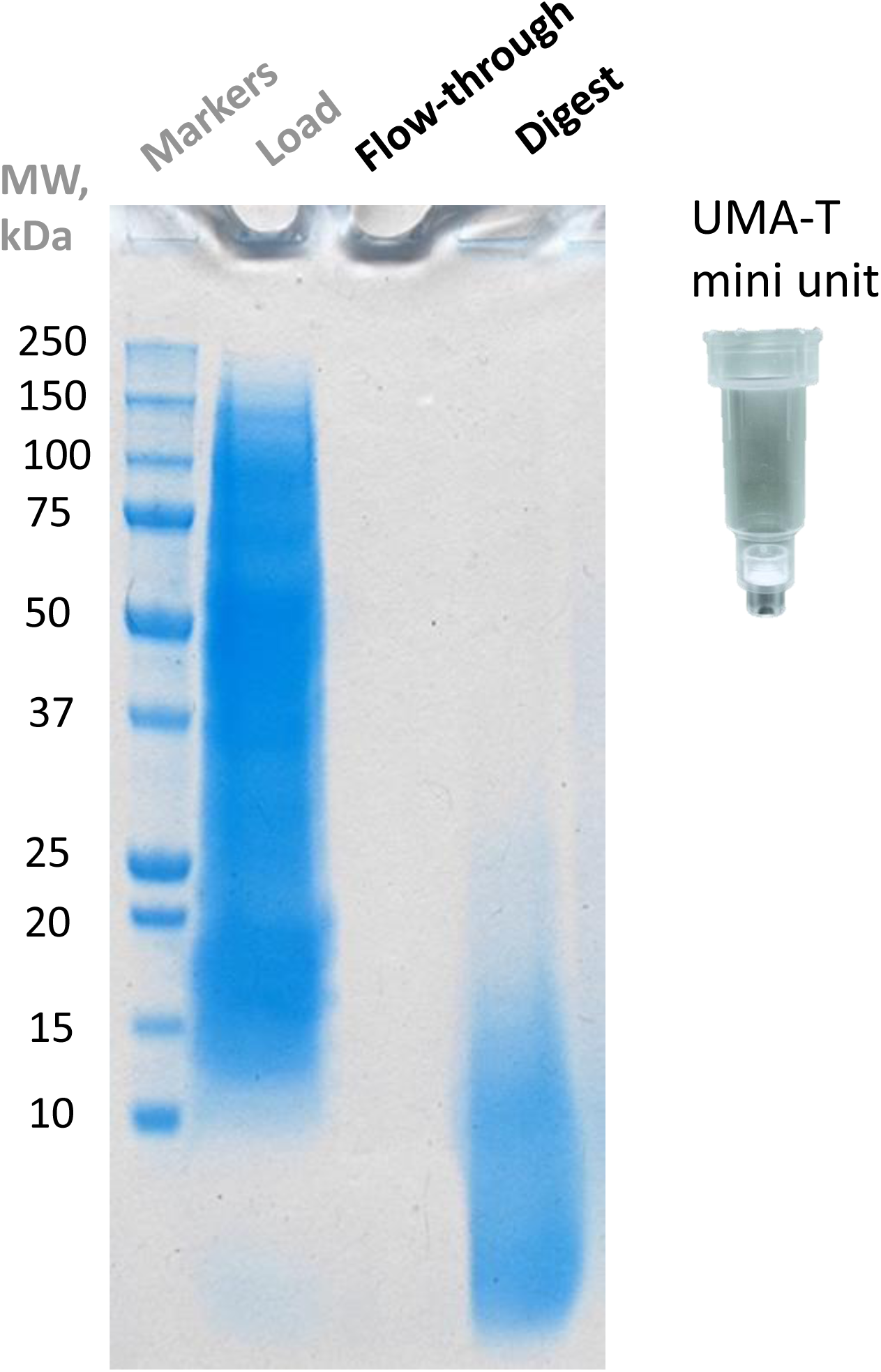
Digestion of HT-29 SDS protein lysate using an UMA paper mini unit with embedded trypsin. Visualization by gel electrophoresis. ∼50 μg of HT-29 cell protein lysate in 4% SDS, 50 mM Tris-HCl, pH 7.6, was processed in the UMA-T embedded trypsin mini unit (the paper-trypsin unit, stored at RT for 2 weeks). Tryptic digest was performed for 45 min at 53 °C in 70 mM AmBic. The digest products were eluted with 70 mM AmBic. The samples were run on a 4-12% Bis-Tris protein gel. The gel was stained with Coomassie.

**Figure 15.**
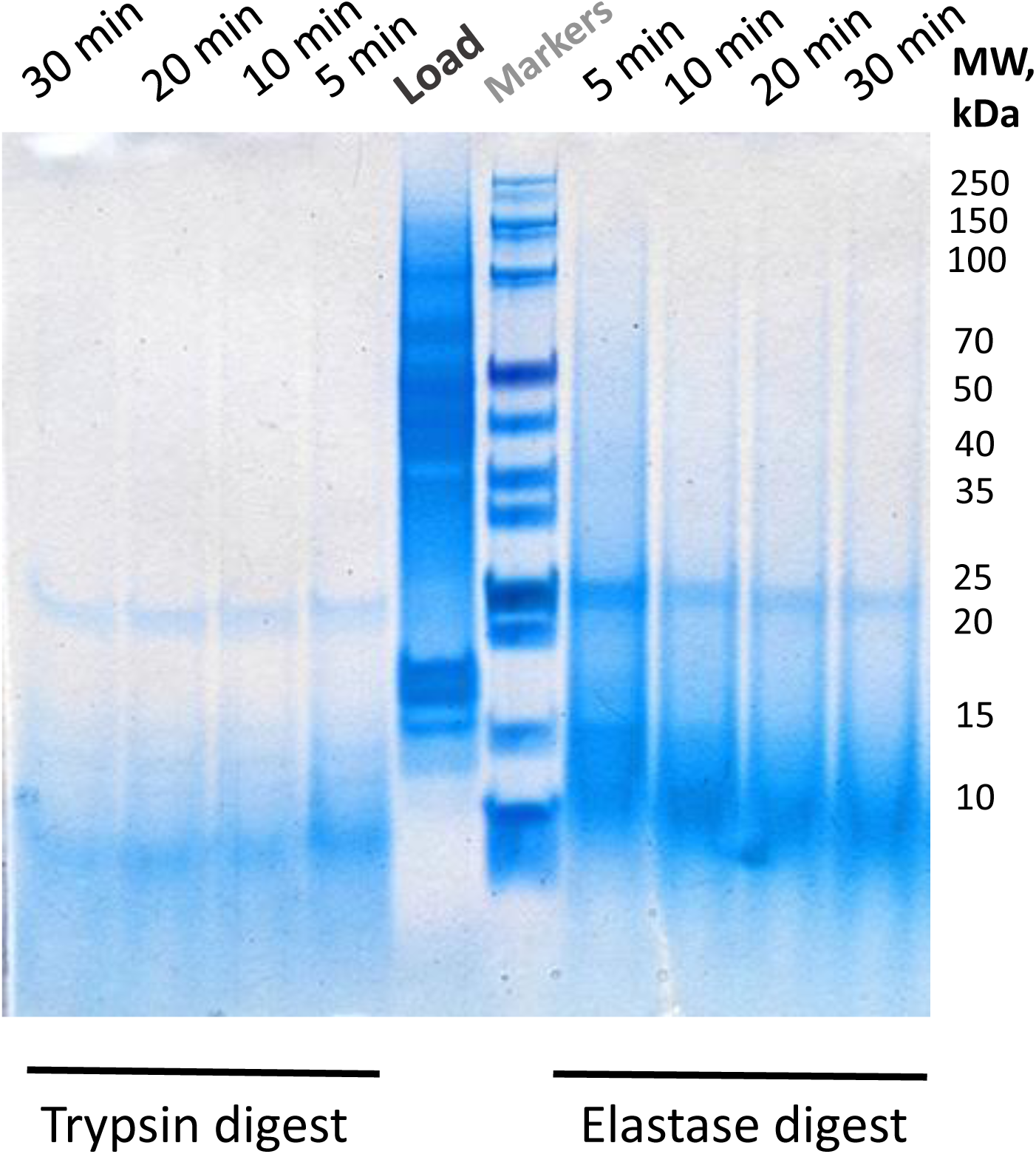
Protein digestion with time intervals of varying lengths using UMA paper tips embedded with trypsin or elastase. UMA paper tips were embedded with either trypsin or elastase. HT-29 SDS lysate was loaded in 6 volumes of the AMA solution. After the wash with the AMA solution, digests at 53 °C in 70 mM AmBic were performed with time intervals of varying lengths (5-30 min). Digest products were eluted with 1X Laemmli buffer. The samples were run on a 4-12% Bis-Tris protein gel. The gel was stained with Coomassie.

As already shown, after the UMA capture protein mixtures could be eluted from the UMA units with trifluoroethanol solutions. The elution with concentrated formic acid is also possible even though there is always the concern regarding the possibility of the associated formylation modifications [12] and practicality using highly concentrated acid solutions.

While working with the remazol-dyed proteins I observed that the cellular lysate proteins solubilized by the basic concentrated trifluoroethanol solution (70% trifluoroethanol, 100 mM triethylammonium bicarbonate) (TT) could be captured on reversed phase media such as wide pore cross-linked polystyrene-divinylbenzene (PS-DVB) Poros R1 and Poros R2 (Thermo) beads which are mostly used in perfusion chromatography [26] (Figure 16). A similar behavior was later observed by using wide pore PS-DVB reversed phase media from other manufacturers - PLRP-S (Agilent) and PolyRP (Sepax). I do not have a plausible explanation for this protein capture phenomenon as the fluorinated alcohol trifluoroethanol has been used comprehensively over the years in polypeptide reversed phase chromatography separations (even though at the acidic pH) both as a mobile phase elution modifier and as a pre-column wash agent to prevent polypeptide carryover between sample injections [22, 27]. The UMA processing unit with incorporated PS-DVB media was called the UMA-R unit. The upper part of this unit, made of either wide microporous filter paper or cotton wool plug, captures the introduced proteins. After the capture and wash, the proteins are forwarded onto and captured by the wide pore (2000 Å – 4000 Å average pore size) reversed phase media bottom part. Given the facts that the average amino acid size is about 3.5 Å, the mean human protein sequence is about 500 amino acids [28], and the proteins are denatured, i.e. partially or completely unfolded, the use of the wide pore perfusion chromatography media (e.g. Poros R1 has an average pore size of 4000 Å, whereas Poros R2 – 2000 Å) for protein recovery and separation is reasonable, in my opinion. In addition, it was found that, similarly to the concentrated FA and basic TFE solutions, the captured proteins could be eluted from the upper part and trapped in the lower RP part of the UMA-R units using 20% Ethanolamine. The captured proteins could then be eluted from the RP material by concentrated acetonitrile or acetonitrile-alcohol mixtures (Supplementary Figures 18 and 19). The findings were confirmed by gel electrophoresis using a cellular protein lysate (Figure 17). It appeared that the use of 20% Ethanolamine resulted in better recovery of low molecular weight proteins as compared with the basic TFE solution. Based on my preliminary observations the UMA-R tips composed of POROS R1 media provide better protein recovery over the UMA-R tips made of PLRP-S or PolyRP media. Additionally, better protein recovery from the UMA-R units could be achieved by using acetonitrile-alcohol elution mixtures, e.g. 50% Acetonitrile, 30% Methanol or 50% Acetonitrile, 30% Isopropanol, in 0.1% trifluoroacetic acid (Supplementary Figures 20 and 21). I think when working with protein load and recovery from the UMA-R units, the use of 20% Ethanolamine for the protein RP loading is preferential as, on one hand, it provides a similar to the concentrated formic acid protein recovery pattern and, on the other hand, in comparison to the concentrated FA, it seems to be less harmful and may carry a lesser chance of the unwanted potential chemical modification of proteins.

**Figure 16.**
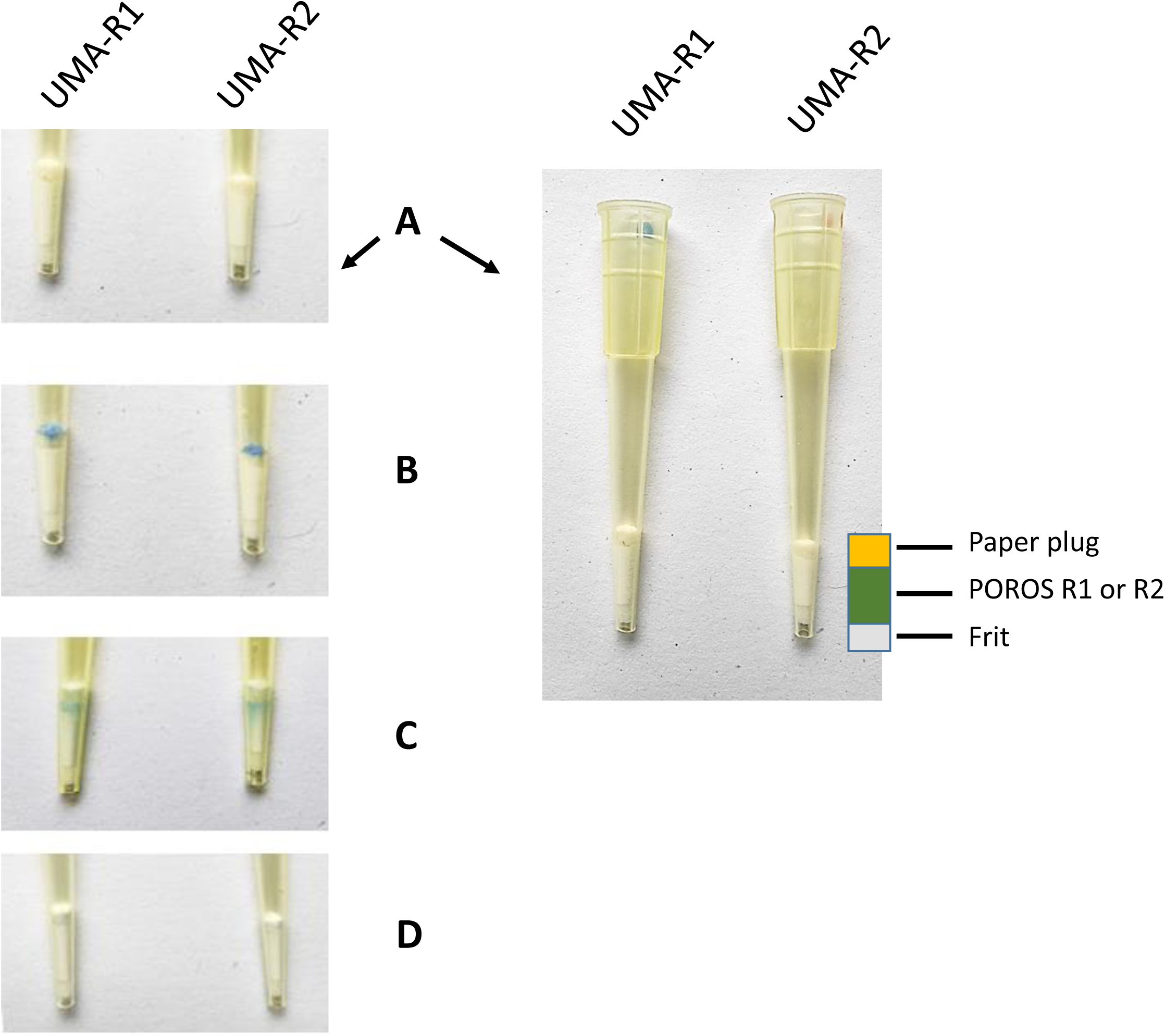
Protein capture and elution using POROS R1 or POROS R2 reversed phase PS-DVB material incorporated into UMA paper tips. Dyed PC3 cell SDS protein lysate was used. UMA-R tips were prepared using either POROS R1 or POROS R2 media (bottom part) and GB003 filter paper plug (upper part) **(A)**. ∼30 μg of PC3-remazol dyed protein SDS lysate was loaded onto the upper paper plug in the AMA solution **(B)**. The proteins were forwarded onto the R1 or R2 tip part with 70% TFE in 100 mM TEAB **(C)**. The proteins were eluted with the 50% Acetonitrile, 30% Isopropanol, 0.1% Trifluoroacetic Acid elution solution **(D)**.

**Figure 17.**
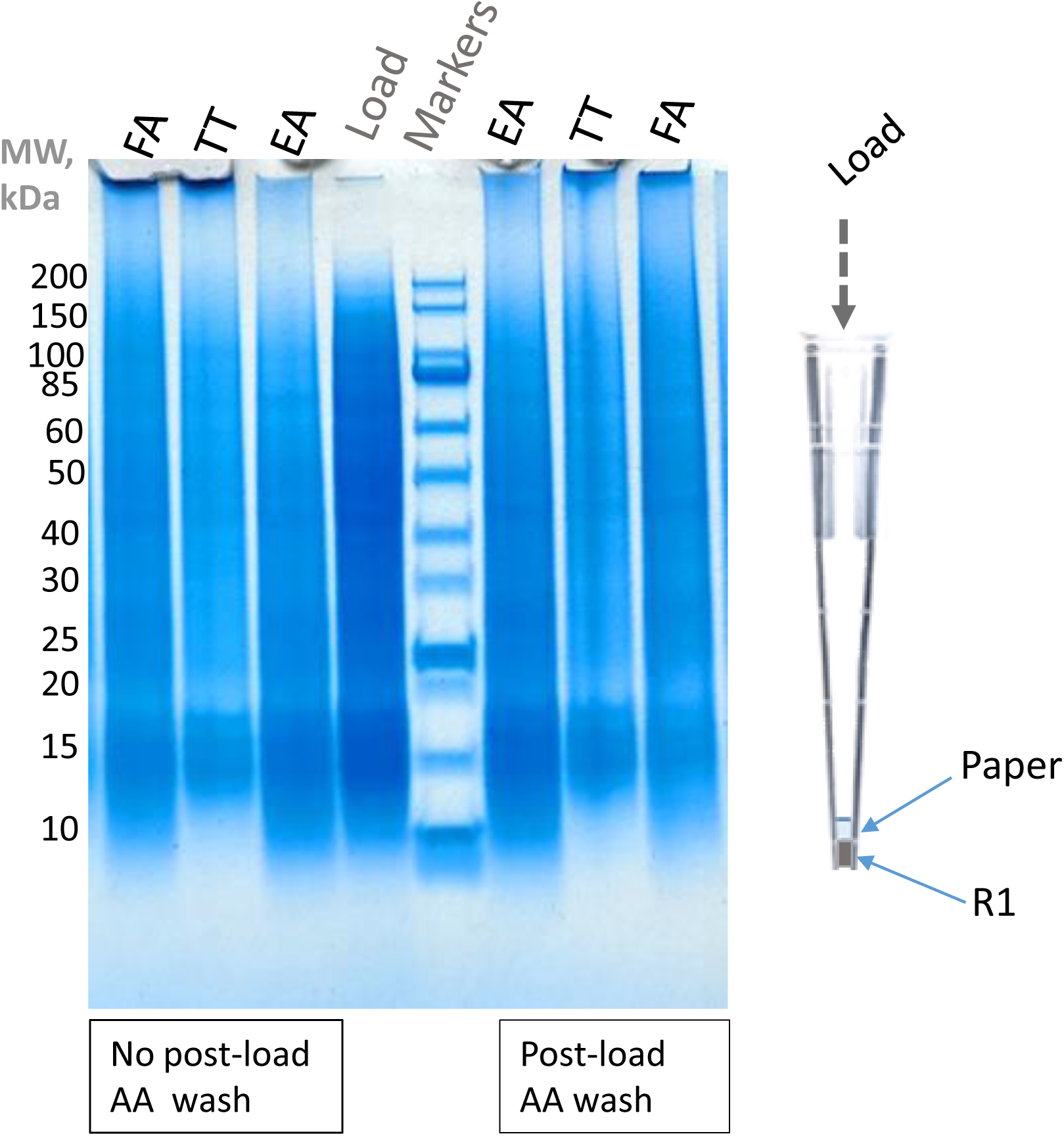
COS-7 cell SDS protein lysate capture and elution from UMA-R tips using different solvents to forward proteins from the upper paper part onto the lower reversed phase part of the UMA-R tip. Visualization by gel electrophoresis. COS-7 cell lysate in 4% SDS was loaded into UMA-R tips (upper part – filter paper, bottom part – POROS R1 media) in 6 volumes of the AMA solution (45% acetonitrile (ACN), 45% methanol, 100 mM Ammonium Acetate (AA)). After the post-load AMA wash either followed or not followed by the wash with 100 mM AA, the paper-captured proteins were eluted onto the R1 reversed phase media after incubation for 2 min at 70 °C with either 20% Ethanolamine (EA), 70% TFE in 100 mM TEAB (TT) or 50% Formic Acid (FA). Following the wash with 3% ACN, the proteins were eluted from the R1 media with 75% ACN in 0.1% TFA. The eluates were dried down and reconstituted with 1X Laemmli buffer. The samples were run on a 4-12% Bis-Tris protein gel. The gel was stained with Coomassie.

I believe that, similarly to the concept described by Bache et al. for peptide separation where the elution from disposable-tip pre-columns was incorporated into the liquid chromatography system operation [29], the concept which has become the basis of the popular Evosep One (Evosep) high throughput reversed phase peptide separation chromatographic platform, the UMA-R unit, with proteins loaded onto its RP part offline, could serve as the disposable pre-column for the consequent chromatographic separation and MS analysis of whole proteins. In addition to the whole protein separation, the UMA-R unit could be used for proteomics experiments involving protein cleavage with proteins trapped in its upper filter part, cleaved, the cleavage products captured in the bottom RP media part and then eluted (Supplementary Figure 21).

The presented Unspecific Molecular Adsorption (UMA) sample preparation methodology provides straightforward means for bottom-up and whole protein analysis. The reported UMA-linked embedded enzyme approach simplifies the routine handling of protein digests, and, together with the described chemical cleavage approach, could become a useful tool for protein sequence characterization. I believe that the observed binding of the enzymes from their diluted solutions in deionized water to the filter paper and other materials is not limited to the tested enzymes, may include additional proteins and find its use in other research areas. The work is in planning to hyphenate the clean-up, concentration and storage of proteins on the UMA-R tips with their consequent chromatographic separation and analysis using up-to-date mass spectrometric systems capable of two-dimensional scanning such as, for example, trapped ion mobility-time-of-flight mass spectrometry [30]. Such a combination, in my view, should facilitate MS profiling of complex whole protein mixtures.

## Associated Data

The mass spectrometry data files were deposited into and are available on MassIVE (ftp://MSV000100759@massive-ftp.ucsd.edu).

## Acknowledgements

The author is grateful to his colleagues at the Biomolecular Mass Spectrometry Facility of the University of Leeds for their support.

A patent application related to this work has been filed.

## Abbreviations

2DE: two-dimensional gel electrophoresis
AA: ammonium acetate
ACN: acetonitrile
AMA: 45% methanol, 45% acetonitrile in 100 mM ammonium acetate
AmBic: ammonium bicarbonate
BUT: butanol
DTT: dithiothreitol
EA: ethanolamine
ESI: electrospray ionization
FA: formic acid
HCl: hydrochloric acid
IAA: iodoacetamide
ISO: isopropanol
LC-MS: liquid chromatography mass spectrometry
LC-MS/MS: liquid chromatography tandem mass spectrometry
MeOH: methanol
MS: mass spectrometry, mass spectrometric
MW: molecular weight
PS-DVB: polystyrene-divinylbenzene
Q-TOF: quadrupole-time-of-flight
RD: remazol dye
RIPA: radioimmunoprecipitation assay buffer
RP: reversed phase
RT: room temperature
SCX: strong cation exchange
SDS: sodium dodecyl sulfate
SPE: solid phase extraction
TCEP: tris(2-carboxyethyl)phosphine hydrochloride
TEAB: triethylammonium bicarbonate
TF: 70% trifluoroethanol, 1% formic acid
TFA: trifluoroacetic acid
TFE: trifluoroethanol
TIC: total ion current
Tris: tris(hydroxymethyl)aminomethane
TT: 70% trifluoroethanol, 100 mM triethylammonium bicarbonate
UMA: unspecific molecular adsorption
UMA-C18: UMA unit incorporating C18 reversed phase material
UMA-T: UMA unit with embedded trypsin
UMA-R: UMA unit incorporating wide pore reversed phase material
UMA-P: UMA pipetting unit

---

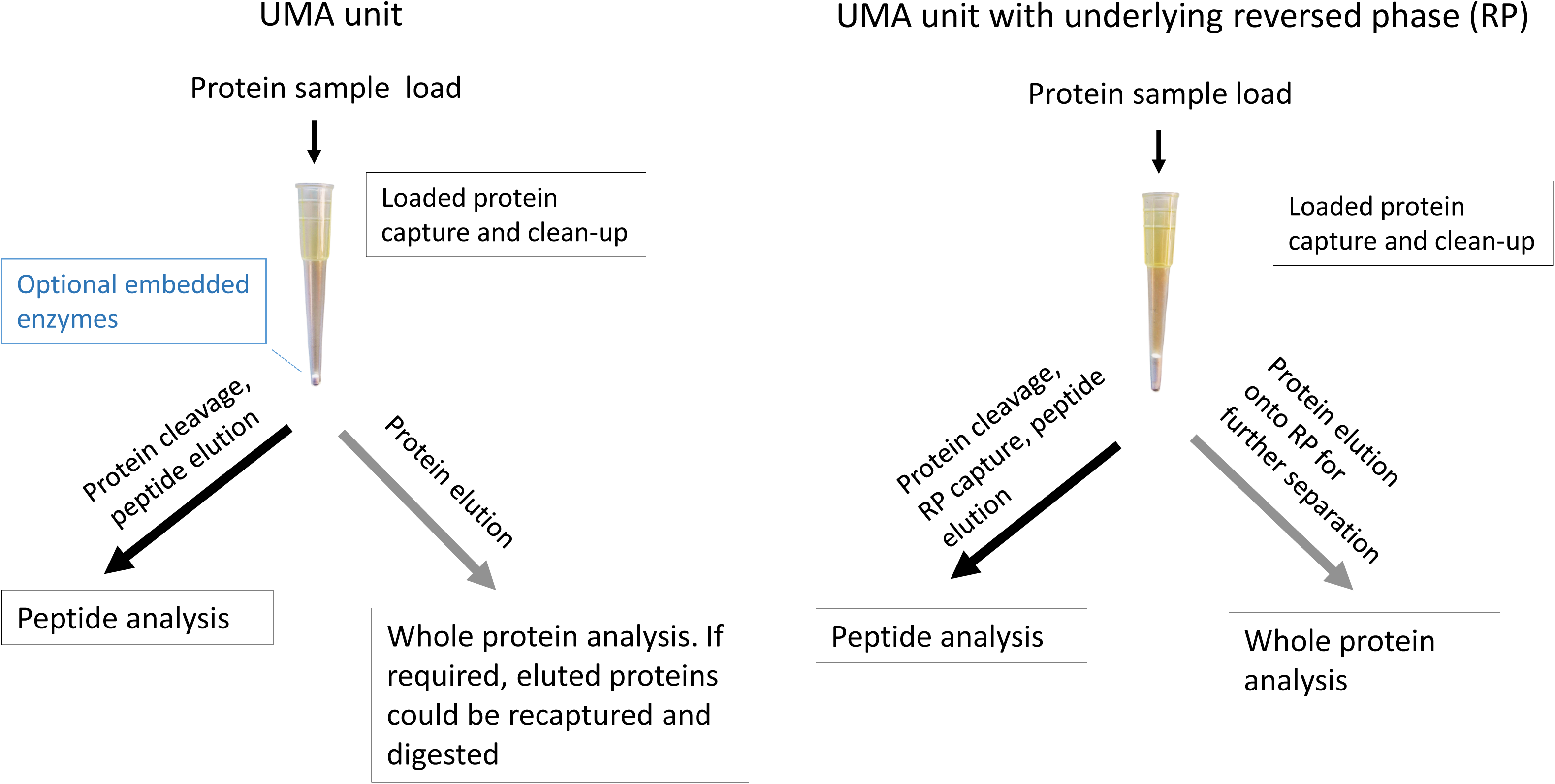
UMA methodology outline (cover figure)

