## Supplementary Figures for "Unspecific Molecular Adsorption (UMA) sample preparation method for bottom-up and whole protein analysis. The foundation"

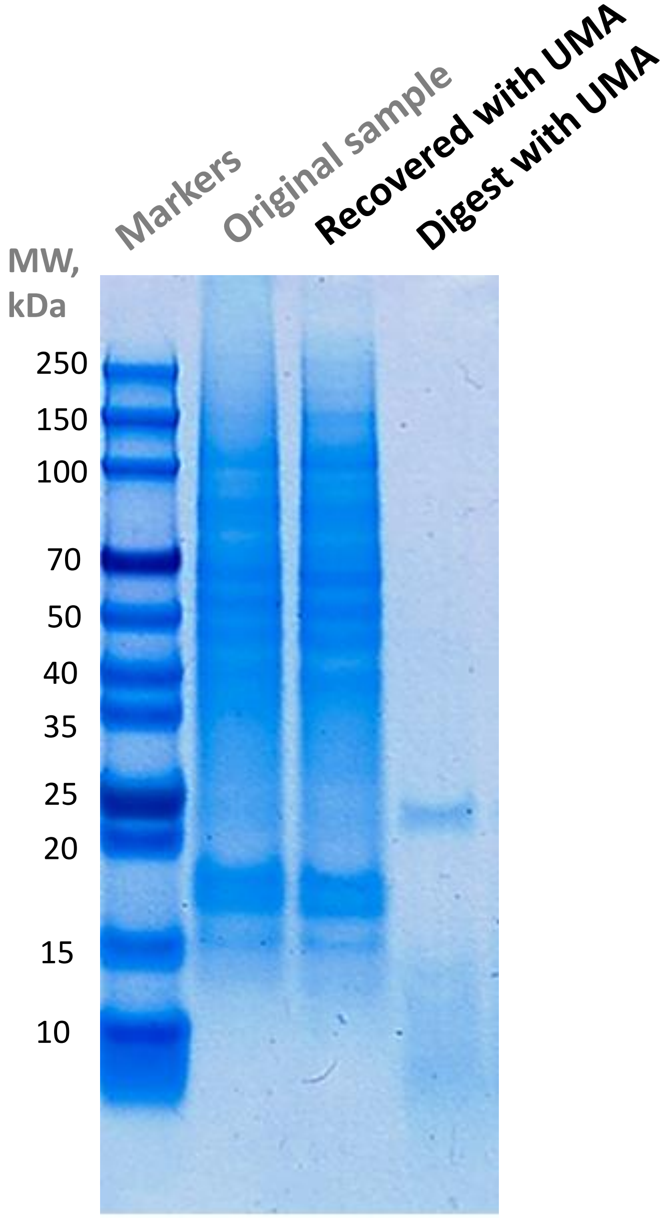


**Capture, recovery and digestion of TFE-extracted proteins on UMA paper mini columns visualized by gel electrophoresis.**

HT-29 cells were lysed with the basic 70% TFE in 100 mM TEAB solution and sonicated. The proteins were further reduced and alkylated with DTT and IAA. The insolubilized material was removed by centrifugation. To the lysate 2.5 volumes of acetonitrile were added. The proteins were captured on UMA paper mini units and then washed with acetonitrile. The proteins were either recovered from the UMA unit with 1X Laemmli buffer or digested on the UMA unit using trypsin in 70 mM Ammonium Bicarbonate for 45 min at 53 °C. The digest products were eluted with 1X Laemmli buffer. The samples were run on a 4-12% Bis-Tris protein gel. The gel was stained with Coomassie.

**Supplementary Figure 1**

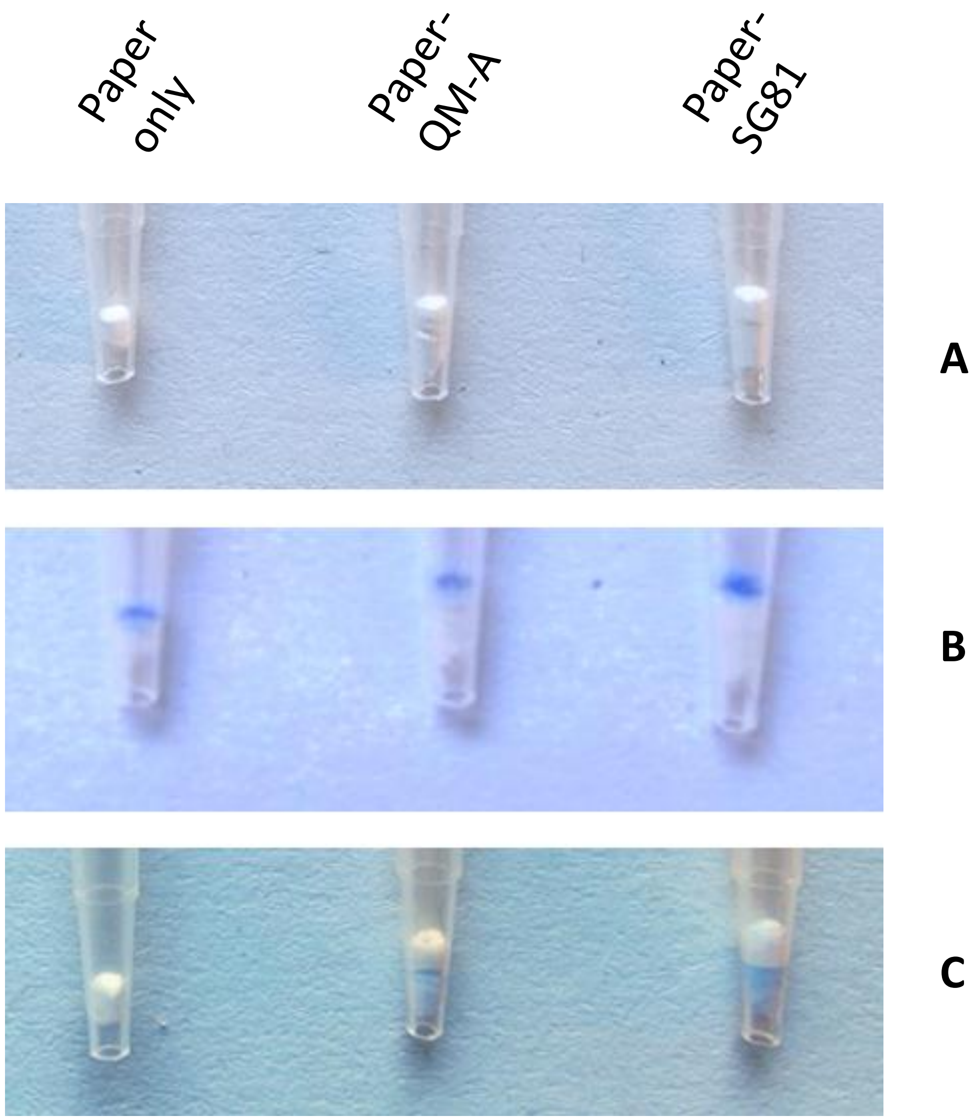


**Testing dyed protein binding to silica under acidic conditions.**
The silica tips were constructed utilizing the GB003 filter paper (upper part) and QM-A quartz silica or SG81 silica paper (bottom part) **(A)**. HT-29 SDS cell protein lysate dyed with Remazol Blue was loaded onto the paper-only, paper-QM-A and paper-SG81 tips with the AMA solution. The tips were washed with the AMA solution and 1% formic acid **(B)**. The proteins were solubilized by the applied 70% trifluoroethanol in 1% formic acid. The solubilized proteins were captured either by the QM-A or SG81 material **(C)**.

**Supplementary Figure 2**

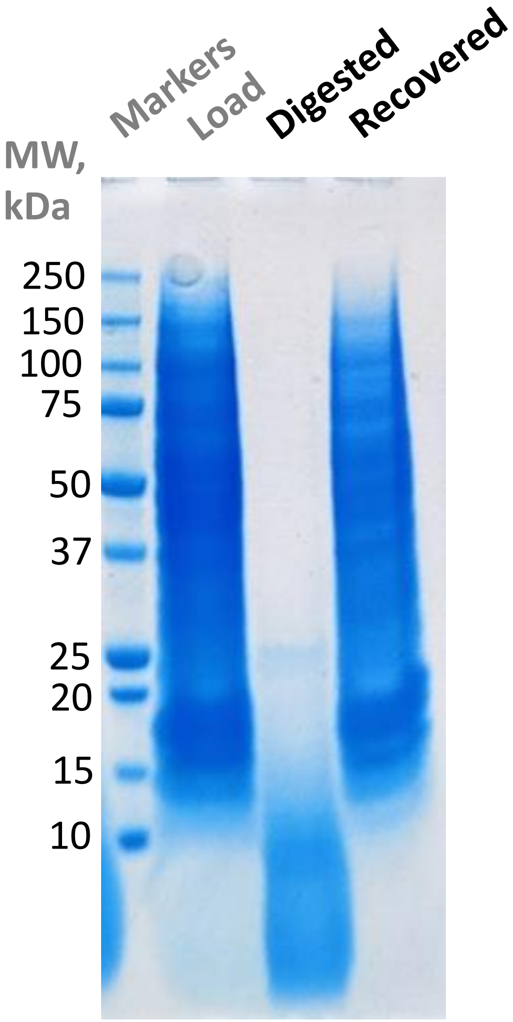


**Testing acidic TFE-extracted protein capture and digestion on SG81 silica paper. Visualization by gel electrophoresis.**
~50 µg of HT-29 cell protein lysate in 4% SDS, 50 mM Tris-HCl, pH 7.6, was captured on UMA paper mini units, solubilized and eluted with 70% TFE in 1% FA and captured on SG81 paper mini units which were prewashed with ACN and 1% FA. The captured material was washed with 70% TFE and then it was either digested with trypsin in 100 mM AmBic and eluted with 1X Laemmli buffer or directly recovered with 1X Laemmli buffer. The samples were run on a 4-12% Bis-Tris protein gel. The gel was stained with Coomassie.

**Supplementary Figure 3**

UMA mini column

UMA tip

UMA pipetting tip

UMA tip with reversed phase

**Examples of the UMA units**

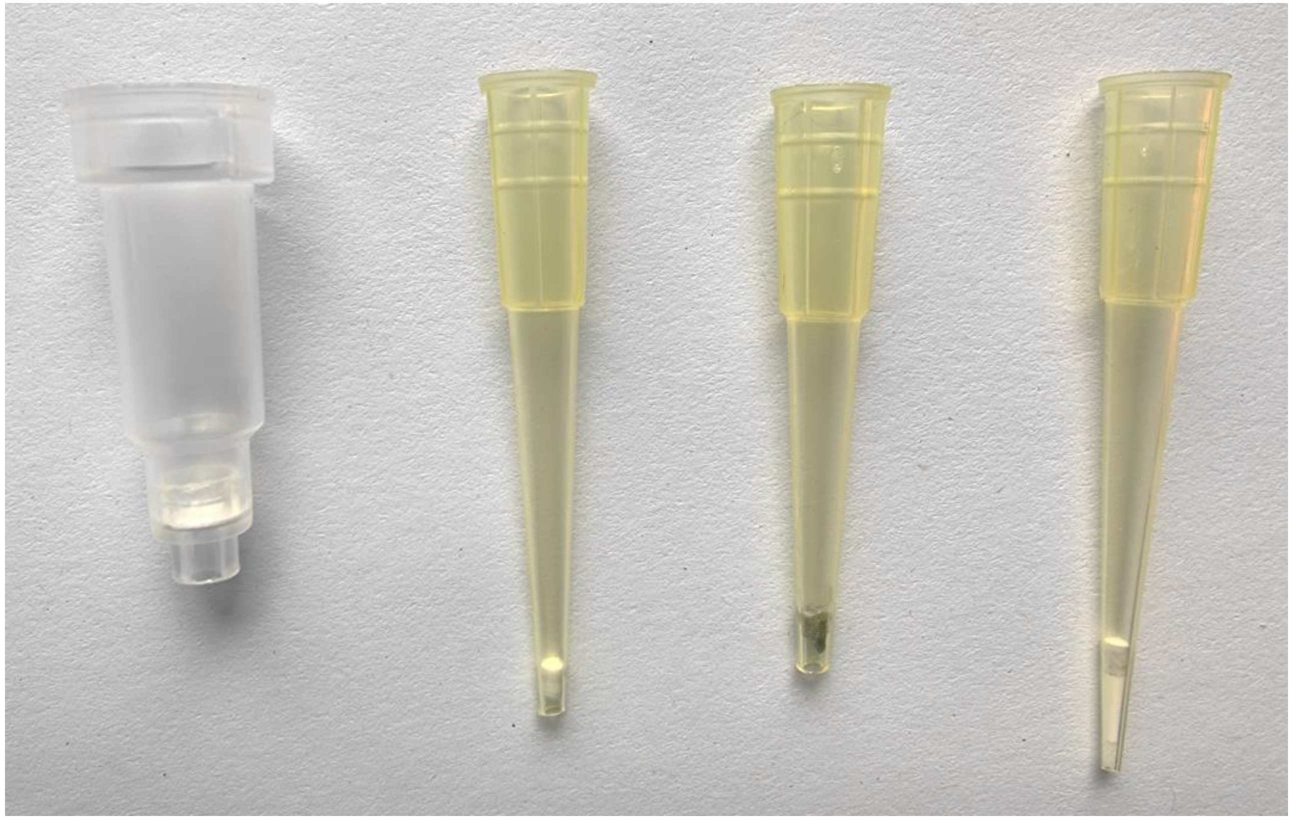

**Supplementary Figure 4**

**Supplementary Figure 5**

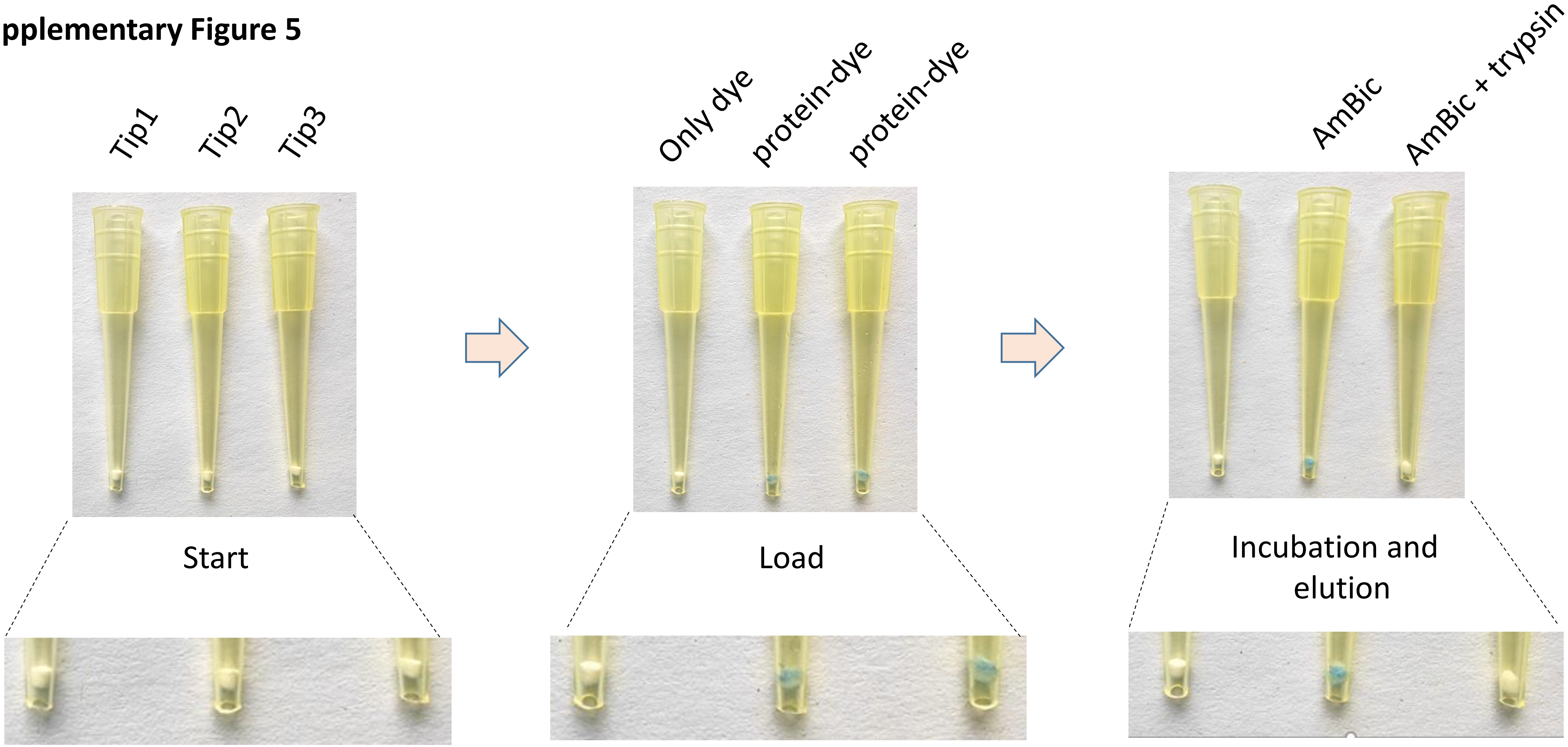


**Protein capture and digestion on UMA paper tips using HT-29 SDS protein lysate dyed with Remazol Blue.**
Using the AMA loading, to the first UMA paper tip Remazol dye in SDS was loaded (no dye capture was observed), to the other two tips ~30 µg of the dyed Remazol HT-29 protein lysate was added (protein-dye capture was observed). One of these two tips was incubated with 70 mM AmBic only, another one was incubated with trypsin in 70 mM AmBic at 56 °C for 15 min. After the incubation, the elution was performed with 70 mM AmBic. The dyed protein elution was observed solely in the enzyme-treated tip.

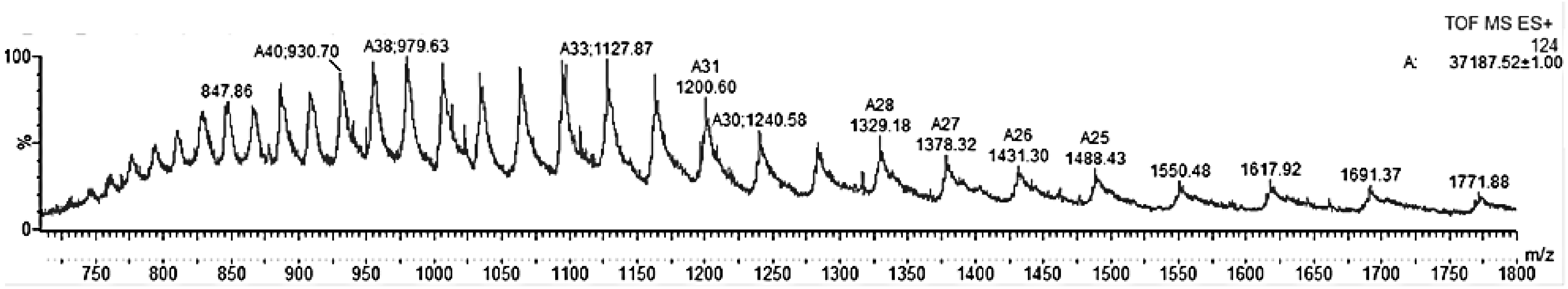


**MS analysis of Alcohol Dehydrogenase solubilized in SDS and cleaned up using an UMA tip with a filter paper plug.**

Alcohol dehydrogenase yeast (A7011, Sigma). Predicted MW 37174.4 Da with N-term Met removed and Cys residues modified with iodoacetamide. The observed delta mass of +13 Da could correspond to an amino acid substitution, e.g. Asp->Gln. The protein was dissolved in 4% SDS, 50 mM TEAB buffer, reduced with DTT and alkylated with iodoacetamide. UMA tip with GB003 paper plug was used. Six volumes of the AMA solution were added to the protein solution aliquot containing 10 µg of protein. The mixture was loaded into the plug by centrifugation. After a wash with the AMA solution, the protein was eluted in 35% TFE, 1% FA. The eluate was analyzed by nanospray in the positive ionization mode on Synapt G1 Q-TOF mass spectrometer.

**Supplementary Figure 6**

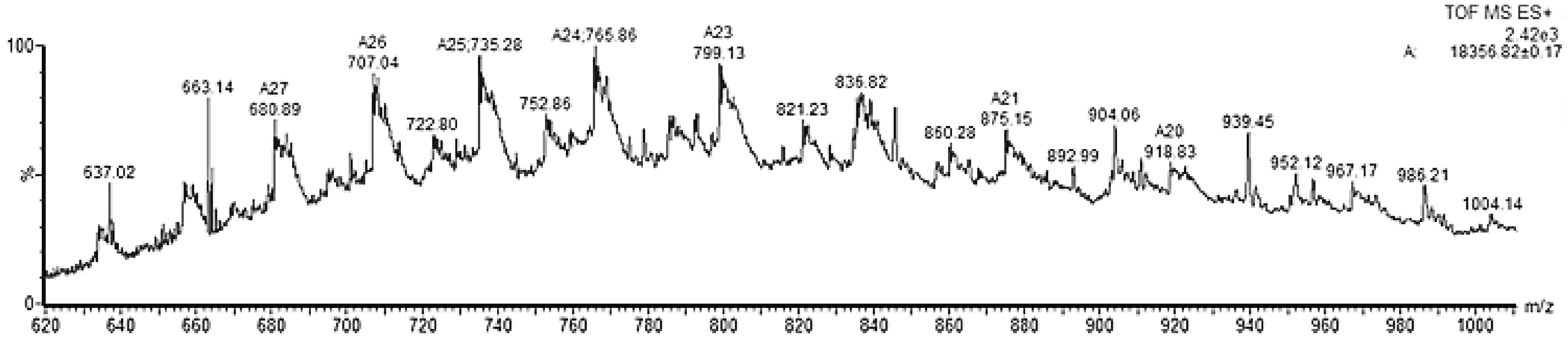


**MS analysis of Myelin Basic Protein solubilized in SDS and cleaned up using an UMA pipetting tip with a felt plug.**

Myelin Basic Protein (Sigma, M1891). Predicted MW 18323.5 Da. The observed delta mass of +33 Da could correspond to oxidation of two Met residues and one deamidation event. The protein mixture was dissolved in 4% SDS, 50 mM TEAB buffer. UMA pipetting tip with a felt plug was used. Five volumes of the AMA solution were added to the protein solution containing 10 µg of the protein. The mixture was loaded into the plug by pipetting up and down several times. After a wash with the AMA, the protein was eluted in 70% TFE, 1% FA. The eluate was analyzed by nanospray in the positive ionization mode on Synapt G1 Q-TOF mass spectrometer.

**Supplementary Figure 7**

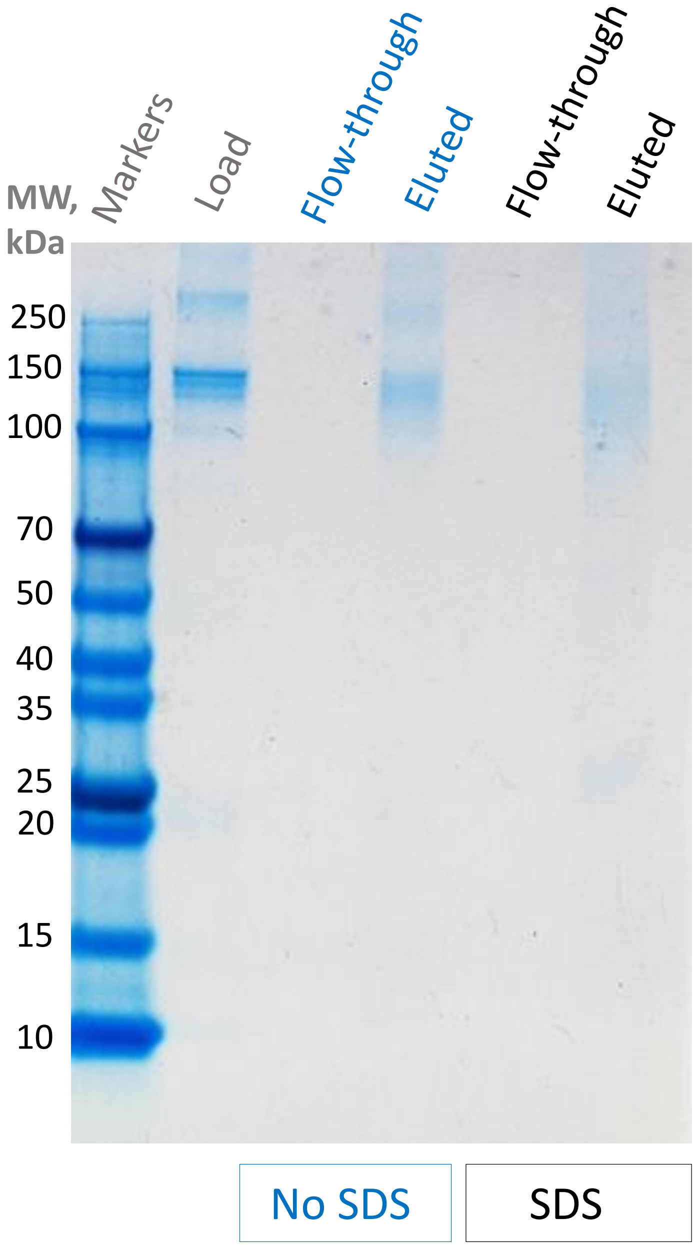


Gel electrophoresis analysis of the eluted antibody

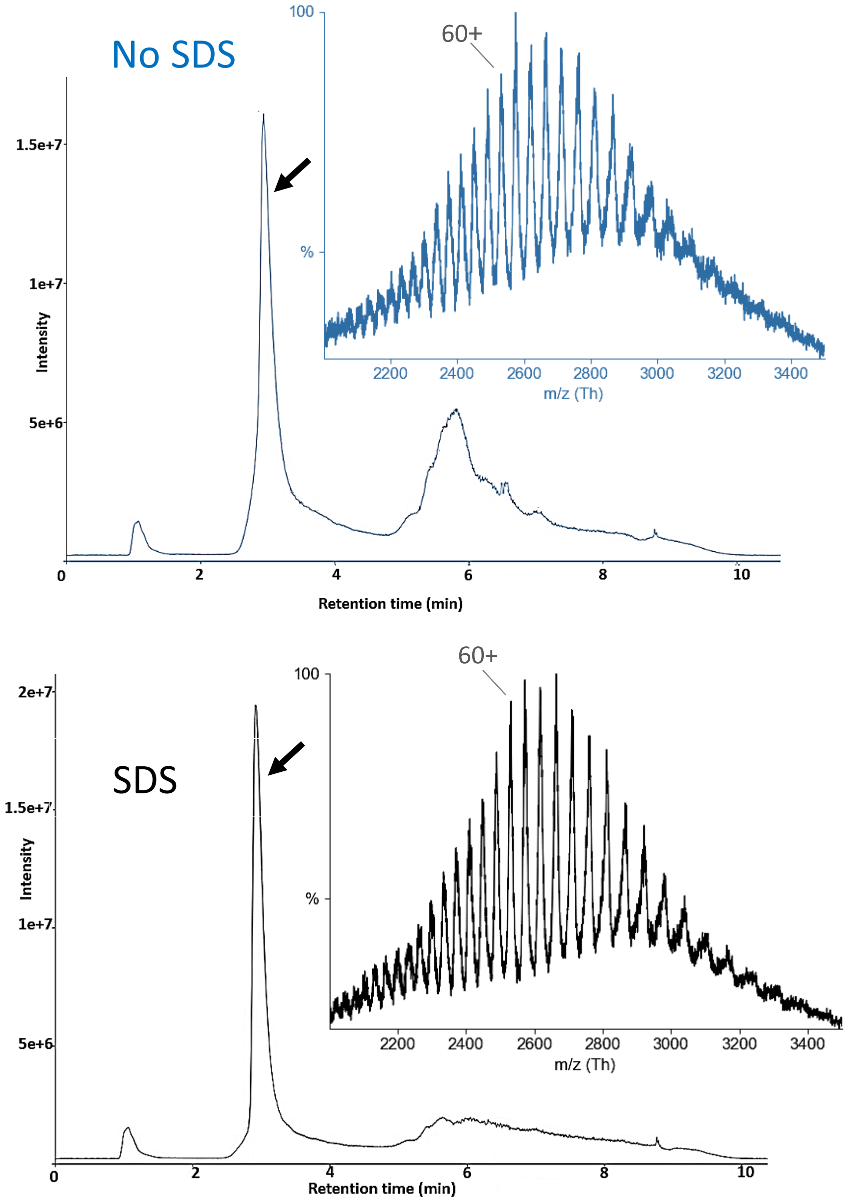


Extracted MS spectrum of the chromatographically separated antibody

**Gel electrophoresis and LC-MS analysis of a monoclonal antibody, with or without SDS, cleaned up using UMA paper tips.**

A commercial monoclonal antibody solution (A2228, Sigma, MW ~150 kDa) was either used as is or had SDS added to it to the final 4% concentration. The antibody samples were loaded onto the UMA paper tips in 6 volumes of the AMA solution, washed with the AMA solution and eluted with 0.1% TFA. For gel electrophoresis, the flow-through and eluate samples were dried down and reconstituted with 1X Laemmli buffer. The samples were run on a 4-12% Bis-Tris protein gel. The gel was stained with Coomassie. In addition, the mass spectrometric profiling of the antibody eluates was done using reversed phase chromatographic separation on the MAbPac™ phenyl column (Thermo) and BioAccord LC-MS TOF System (Waters).

**Supplementary Figure 8**

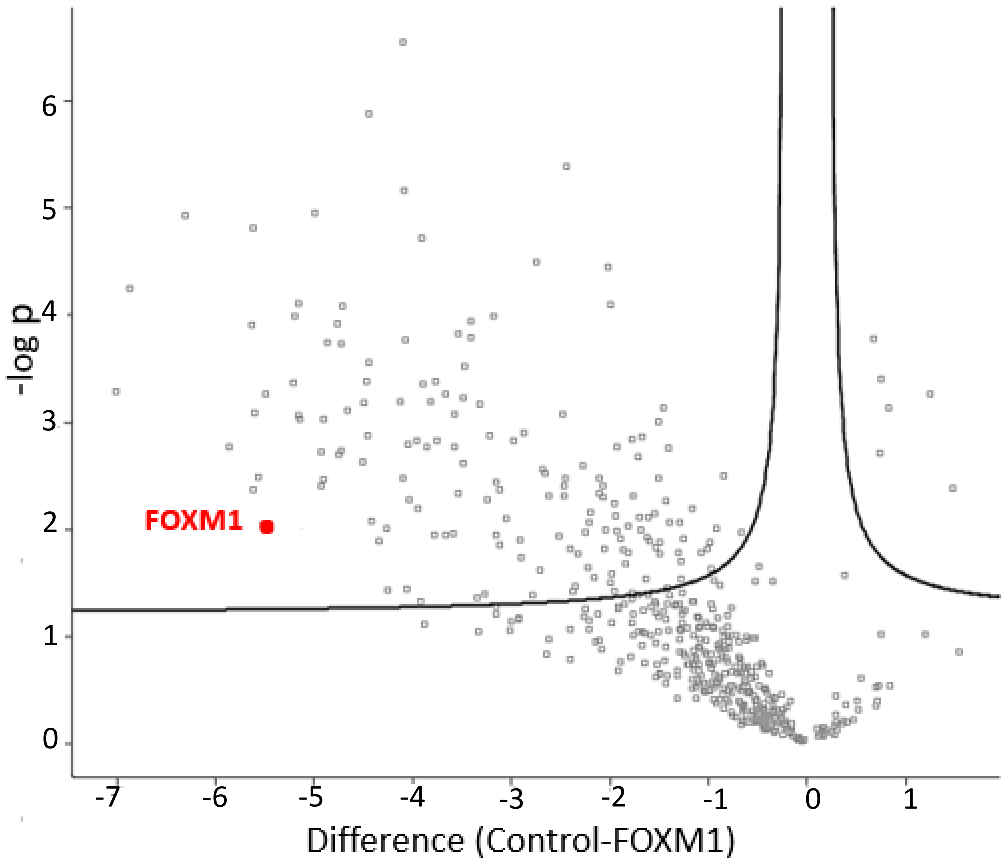


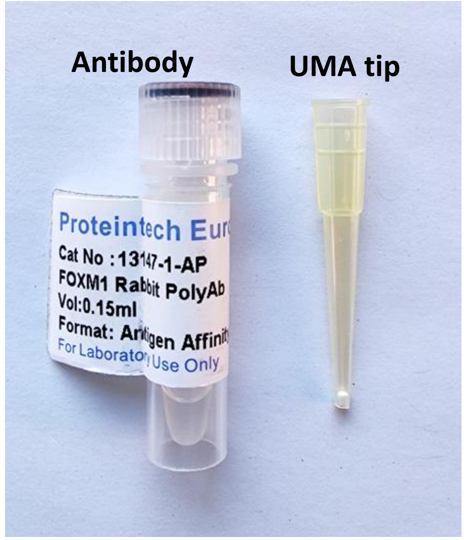


**Analysis of the FOXM1 pull-down experiment from HeLa cells using UMA paper tips.**

To check viability of the rabbit anti-FOXM1 antibody after extended long-term storage (~10 years at 4°C) an immunoprecipitation experiment was performed using 2 µg of this antibody and a control rabbit antibody alongside with 1 mg of HeLa protein lysate, in triplicates. Bound proteins were eluted with 4% SDS and processed using UMA paper tips with digestion by trypsin. The resultant peptides were run on LC-MS Exploris 240 system. Proteins were plotted using Perseus (https://maxquant.net/perseus/) by fold change (Difference) and significance (-log p) using false discovery rate (FDR) of 0.05 and S0 of 0.1. FOXM1 was clearly identifiable in the tested anti-FOXM1 immunoprecipitates.

**Supplementary Figure 9**

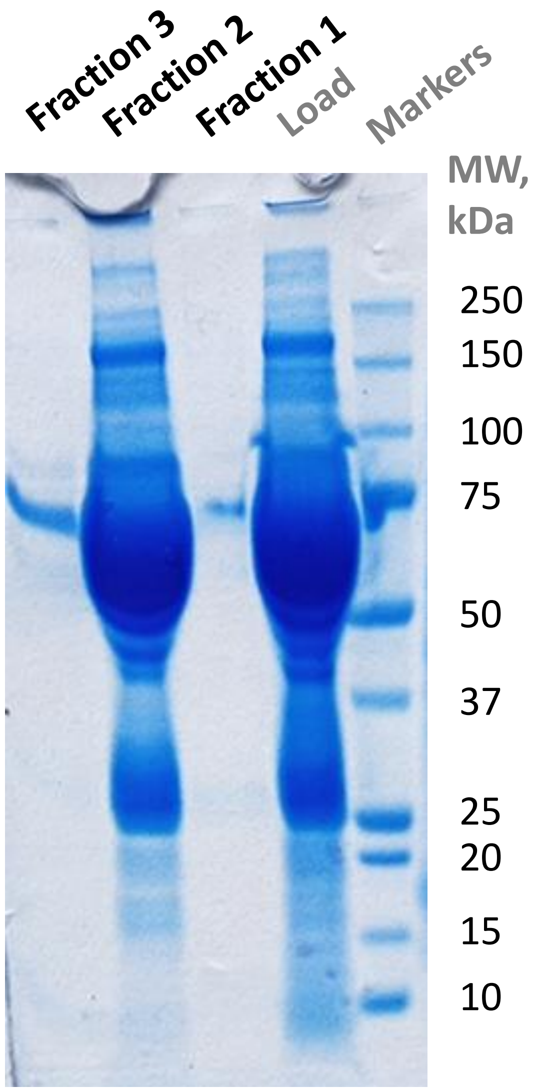


**Fractionation of serum using UMA pipetting units. Visualization by gel electrophoresis.**
1 µl of serum was diluted in 100 µl of the 250 mM TEAB, 5 mM TCEP, 10 mM IAA buffer solution and incubated at RT for 10 min. The mixture was pipetted through an UMA-P (stainless steel mesh) tip several times which was followed by the wash with 250 mM TEAB (fraction 1). Then, 2 volumes of ACN were added to the mixture and it was pipetted up and down again using a new UMA-P tip which was followed by the wash with 70% ACN, 80 mM TEAB (fraction 2). Then, another volume of ACN was added and the mixture was pipetted again using a new UMA-P tip (fraction 3). The bound proteins were eluted from the tips with 1X Laemmli buffer. The samples were run on a 4-12% Bis-Tris protein gel. The gel was stained with Coomassie.

**Supplementary Figure 10**

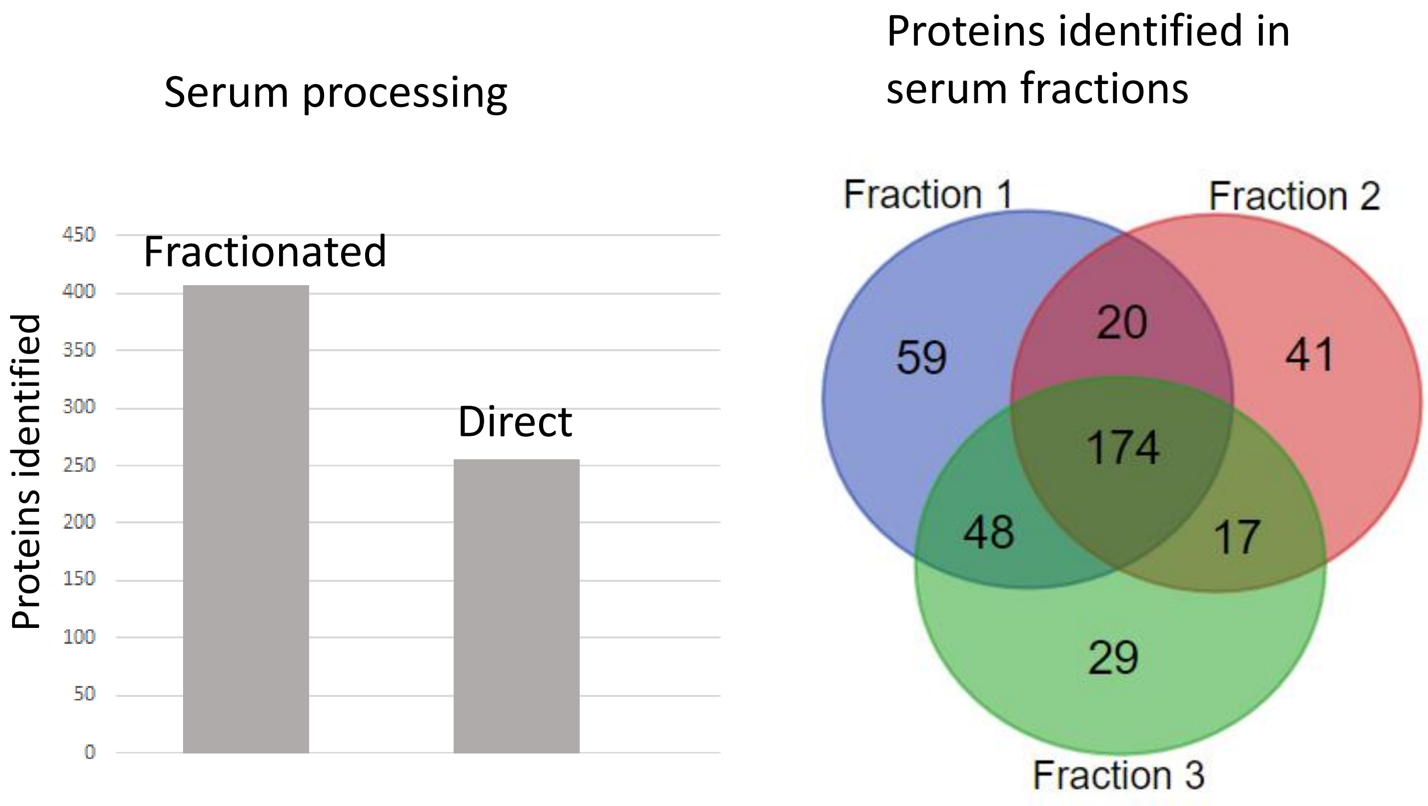


**LC-MS/MS comparison of serum fractionation on UMA pipetting units with direct serum processing on UMA paper units.**
Serum was processed either by fractionation on UMA-P pipetting tips or by direct UMA loading using trypsin digestion for 45 min at 53 °C, in three replicates. The resultant protein digests were run twice using LC-MS Exploris 240 system. The total acquisition time for each run was 50 min. The data were processed by MaxQuant (https://maxquant.net/). For protein identification at least one unique peptide per protein was required.

**Supplementary Figure 11**

**Examples of the ladder-like protein cleavage patterns by elastase and HCl-TCEP.**
Both elastase digestion and acidic cleavage resulted in ladder-like unspecific protein cleavage patterns.

| | Elastase | HCl-TCEP |
|---|---|---|
| Insulin | A.FVNQHL.C<br>A.FVNQHLC.G<br>A.FVNQHLCG.S<br>A.FVNQHLCGS.H<br>A.FVNQHLCGSH.L<br>A.FVNQHLCGSHL.V<br>A.FVNQHLCGSHLVE.A | C.GSHLVEA.L<br>C.GSHLVEAL.Y<br>C.GSHLVEALY.L<br>C.GSHLVEALYL.V<br>C.GSHLVEALYLV.C<br>C.GSHLVEALYLVCG.E<br>C.GSHLVEALYLVCGE.R<br>C.GSHLVEALYLVCGER.G<br>C.GSHLVEALYLVCGERG.F |
| Trypsin | I.SGWGNTK.S<br>I.SGWGNTKS.S<br>I.SGWGNTKSS.G<br>I.SGWGNTKSSGS.S<br>I.SGWGNTKSSGSSYPS.L<br>I.SGWGNTKSSGSSYPSLL.Q<br>I.SGWGNTKSSGSSYPSLLQ.C<br>I.SGWGNTKSSGSSYPSLLQCL.K | N.KPGVYTKVC.N<br>N.KPGVYTKVCN.Y<br>N.KPGVYTKVCNY.V<br>N.KPGVYTKVCNYV.N<br>N.KPGVYTKVCNYVN.W<br>N.KPGVYTKVCNYVNW.I<br>N.KPGVYTKVCNYVNWIQ.Q |
| Carbonic Anhydrase | M.SHHWGY.G<br>M.SHHWGYG.K<br>M.SHHWGYGK.H<br>M.SHHWGYGKH.N<br>M.SHHWGYGKHN.G<br>M.SHHWGYGKHNG.P<br>M.SHHWGYGKHNGPE.H<br>M.SHHWGYGKHNGPEH.W<br>M.SHHWGYGKHNGPEHWH.K<br>M.SHHWGYGKHNGPEHWHK.D<br>M.SHHWGYGKHNGPEHWHKD.F<br>M.SHHWGYGKHNGPEHWHKDFPI.A | G.TYRLVQ.F<br>G.TYRLVQF.H<br>G.TYRLVQFH.F<br>G.TYRLVQFHF.H<br>G.TYRLVQFHFH.W<br>G.TYRLVQFHFHW.G<br>G.TYRLVQFHFHWG.S |
| Serum Albumin | A.KYICEN.Q<br>A.KYICENQ.D<br>A.KYICENQD.S<br>A.KYICENQDS.I<br>A.KYICENQDSI.S<br>A.KYICENQDSIS.S<br>A.KYICENQDSISS.K<br>A.KYICENQDSISSK.L<br>A.KYICENQDSISSKL.K<br>A.KYICENQDSISSKLK.E | K.SEVAHRF.K<br>K.SEVAHRFK.D<br>K.SEVAHRFKDL.G<br>K.SEVAHRFKDLG.E<br>K.SEVAHRFKDLGE.E<br>K.SEVAHRFKDLGEE.N<br>K.SEVAHRFKDLGEEN.F<br>K.SEVAHRFKDLGEENF.K<br>K.SEVAHRFKDLGEENFKA.L<br>K.SEVAHRFKDLGEENFKAL.V<br>K.SEVAHRFKDLGEENFKALVL.I<br>K.SEVAHRFKDLGEENFKALVLI.A |

**Supplementary Figure 12**

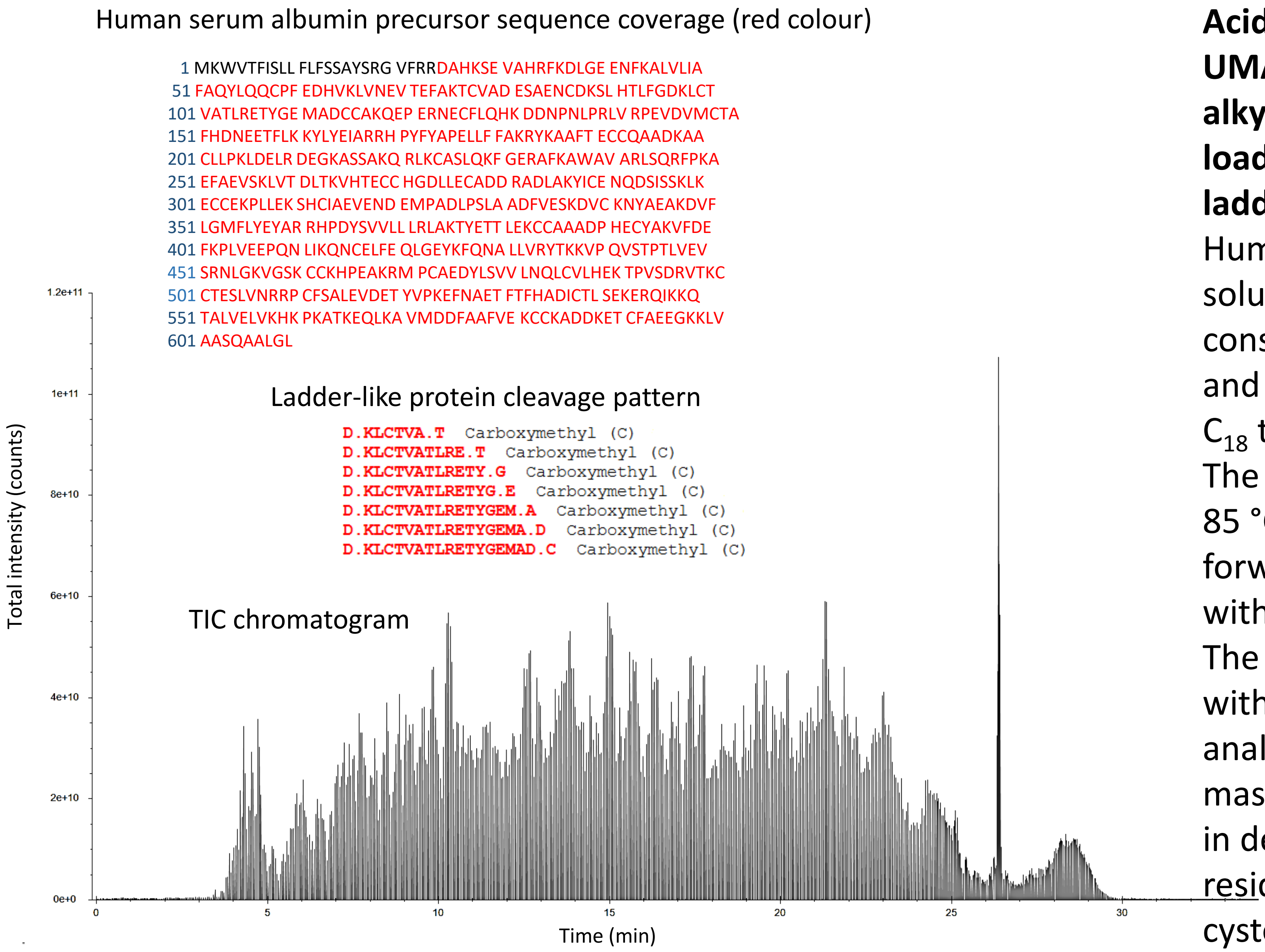


**Supplementary Figure 13**

**Acidic cleavage of human serum albumin on an UMA-$C_{18}$ tip in 10% HCl. The protein was alkylated with iodoacetamide prior to the tip loading. LC-MS/MS sequence coverage and the ladder-like protein cleavage pattern example.** Human serum albumin (A3782, Sigma) was solubilized in the 4% SDS, 50 mM TEAB buffer, consecutively reduced and alkylated with DTT and IAA. The protein was loaded into the UMA-$C_{18}$ tip with six volumes of the AMA solution. The captured protein was cleaved in 10% HCl at 85 °C for 30 min, the peptide products were forwarded onto the $C_{18}$ part of the tip, washed with water and eluted with 60% ACN in 0.2% FA. The eluate was dried down and reconstituted with 1% ACN, 0.2% FA. The peptides were analysed by LC-MS/MS on LTQ-Orbitrap Velos mass spectrometer. The acidic cleavage results in deamidation of asparagine and glutamine residues as well as of carboxyamidomethyl cysteine residues (when protein's cysteine residues were alkylated with IAA prior to the acidic cleavage).

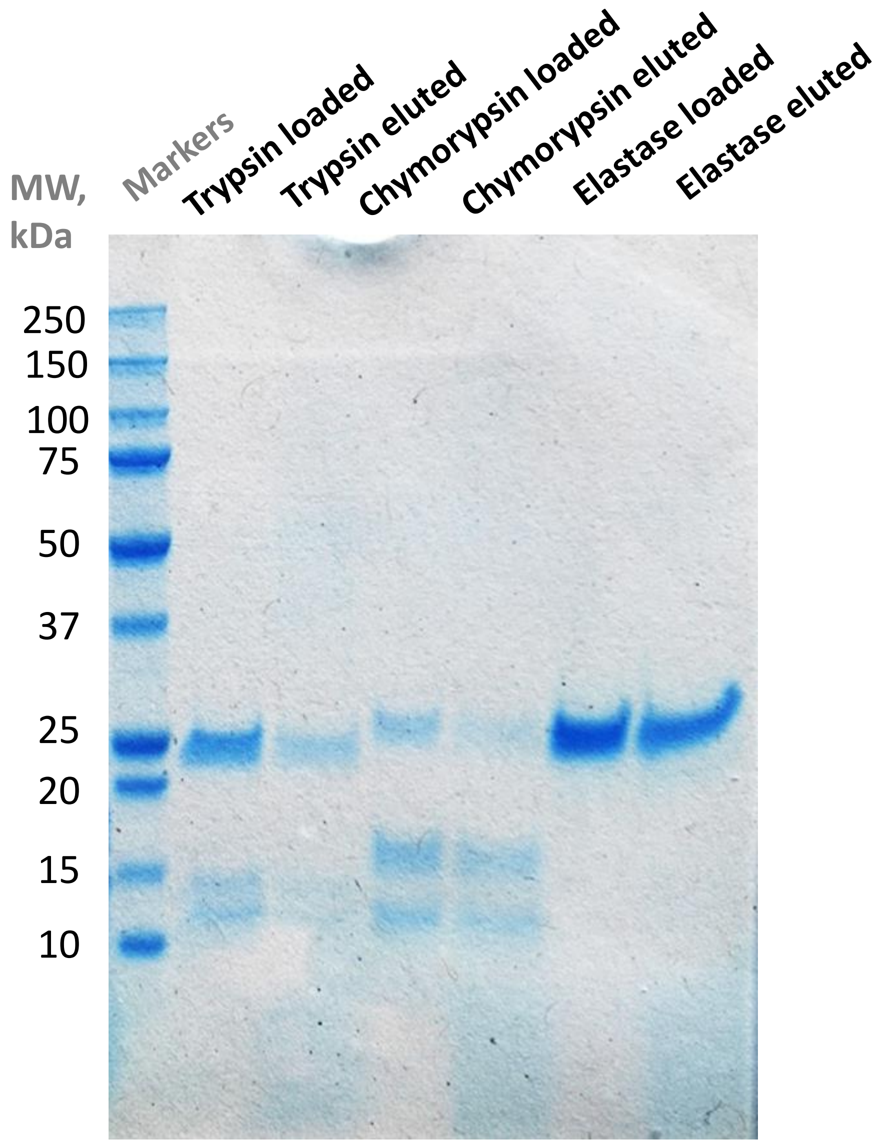


**Capture of enzymes dissolved in deionized water on cotton wool.**
~2.5 µg of trypsin, chymotrypsin and elastase in 200 µl of deionized water were slowly pushed through the UMA tips incorporating cotton wool plugs. The tips then were washed with deionized water and ethanol. The captured material was eluted with 1X Laemmli buffer. The samples were run on a 4-12% Bis-Tris protein gel. The gel was stained with Coomassie.

**Supplementary Figure 14**

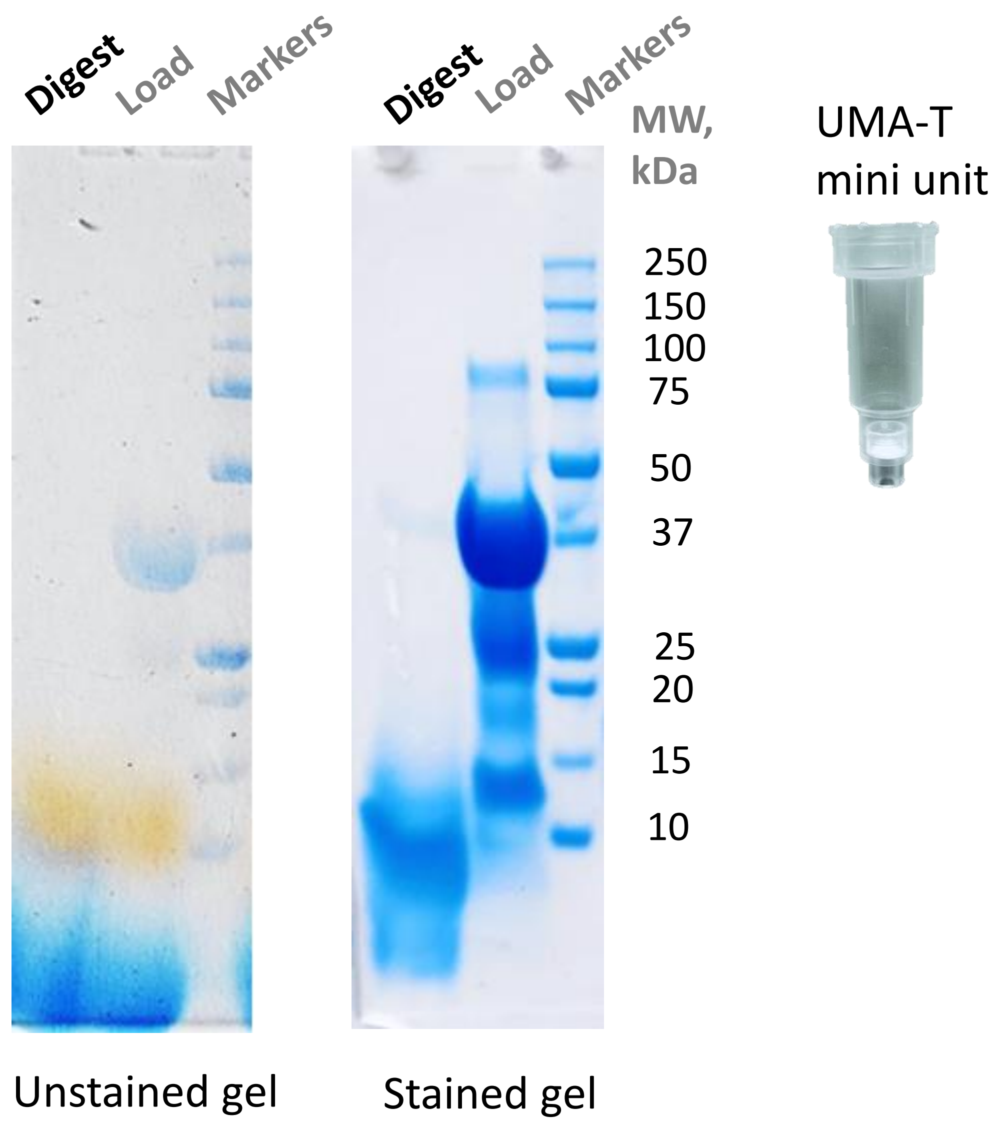


**Digestion of dyed Alcohol Dehydrogenase using an UMA paper mini unit with embedded trypsin. Visualization by gel electrophoresis.**

Alcohol Dehydrogenase (ADH) was dissolved in 4% SDS, 50 mM NaHCO3 and labelled with Remazol Blue (RD) dye (weight ratio ADH:RD = 8:1) overnight at 55 °C. ~30 µg of the dyed product was processed by the UMA-T mini unit (paper-trypsin unit, stored at RT for 2 weeks). The digest was performed for 45 min at 53 °C in 70 mM Ammonium Bicarbonate. The digest products were eluted with 1X Laemmli buffer. The samples were run on a 4-12% Bis-Tris protein gel. The gel was stained with Coomassie.

**Supplementary Figure 15**

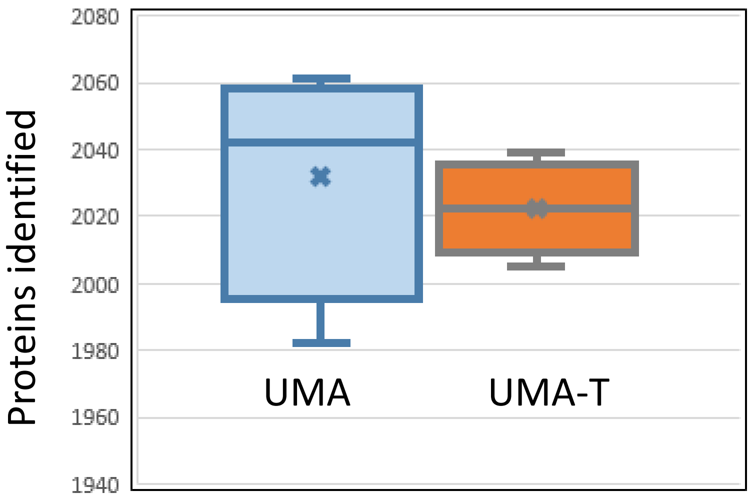


**LC-MS/MS comparison of PC3 cell SDS protein lysate digestion using UMA paper mini units with or without embedded trypsin.**
PC3 cellular protein lysate in SDS was processed by UMA and UMA-T units using either loaded (UMA) or embedded trypsin (UMA-T) digestion in 70 mM AmBic for 45 min at 53 °C in four replicates. The digest products were analyzed by LC-MS/MS using Exploris 240 orbitrap system. The total acquisition time for each run was 45 min. The data were processed by MaxQuant. For protein identification at least one unique peptide per protein was required.

**Supplementary Figure 16**

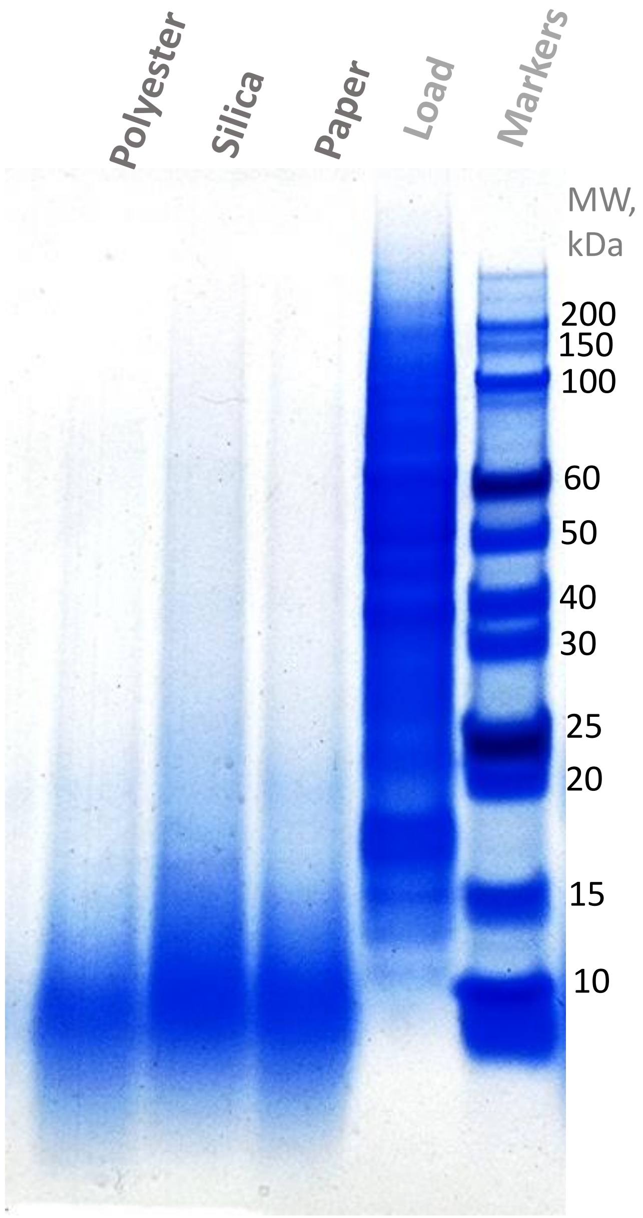


**Digestion of MGH-U3 SDS protein lysate by UMA units made with different materials and embedded trypsin. Visualization by gel electrophoresis.**

Paper-trypsin, silica-trypsin and polyester-trypsin UMA tips were created by passing 2.5 µg of trypsin diluted in deionized water through 200-µl tips incorporating 2-mm paper, silica or polyester plugs. Afterwards, the tips were washed with ethanol. ~50 µg of MGH-U3 cell protein lysate in 4% SDS, 50 mM Tris-HCl, pH 7.6, was processed in these embedded trypsin units using the UMA approach. Digestion was performed for 30 min at 53 °C in 70 mM AmBic. The digest products were eluted with 1X Laemmli buffer. The samples were run on a 4-12% Bis-Tris protein gel. The gel was stained with Coomassie.

**Supplementary Figure 17**

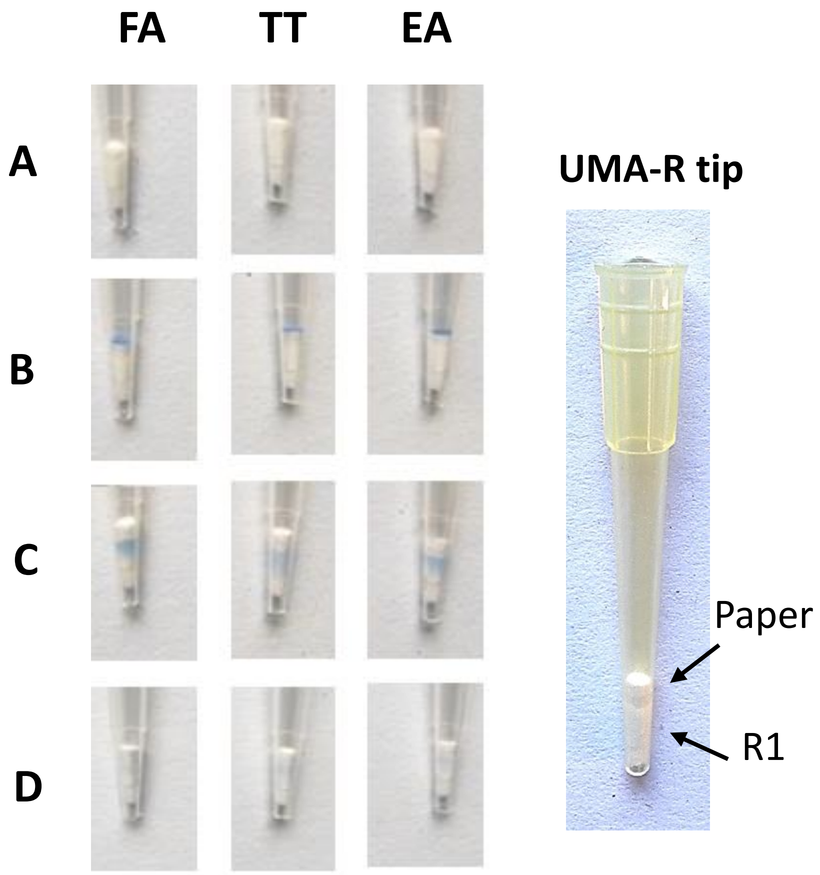


**Dyed HT-29 cell SDS protein lysate capture and elution from UMA-R tips which incorporated POROS R1 material into UMA paper tips.** UMA-R tips were constructed using the GB003 filter paper (upper part) and POROS R1 PS-DVB materials (bottom part) **(A)**. HT-29 SDS cell protein lysate dyed with Remazol Blue was loaded onto the tips with the AMA solution **(B)**. The paper-captured proteins were transferred onto the R1 part using one of the following solutions: 20% Ethanolamine (**EA**), 70% TFE in 100 mM TEAB (**TT)** and 50% Formic Acid (**FA**) **(C)**. After the R1 protein capture, proteins were eluted with 75% Acetonitrile in 0.1% TFA **(D).**

**Supplementary Figure 18**

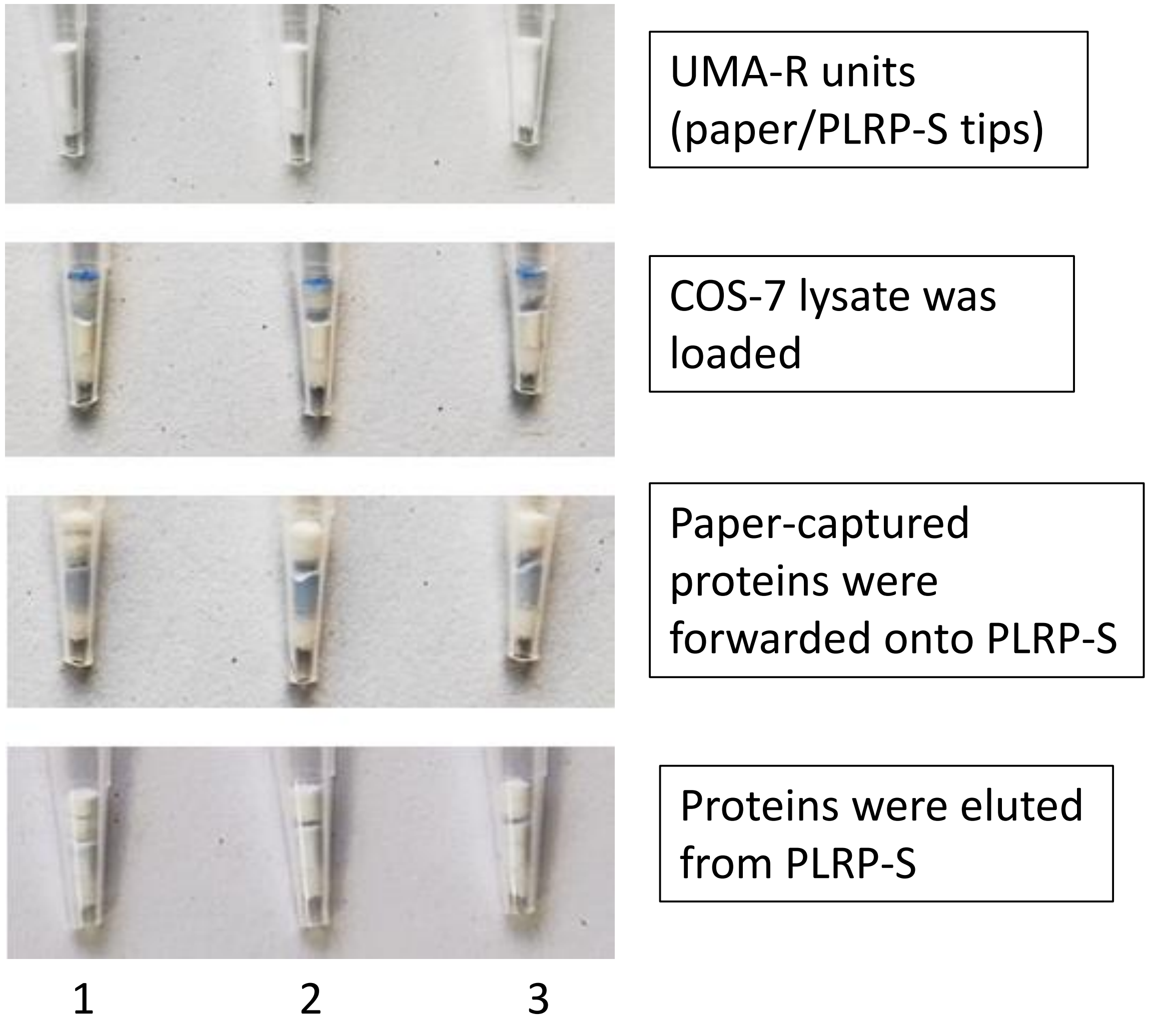


**Dyed COS-7 cell SDS protein lysate capture and elution from UMA-R tips which incorporated PLRP-S reversed phase material into UMA paper tips.** UMA-R tips contained the upper GB003 paper part and bottom PLRP-S media part. COS-7 cell lysate in 4% SDS dyed with Remazol Blue was loaded onto the UMA-R tips in 6 volumes of the AMA solution (45% acetonitrile, 45% methanol, 100 mM Ammonium Acetate (AA)). After the post-load AMA wash, followed by the wash with 100 mM AA, the paper-captured proteins were eluted onto the PLRP-S reversed phase media after incubation for 2 min at 70 °C with 20% Ethanolamine. Following the wash with 3% acetonitrile, the proteins were eluted from the PLRP-S media with different combinations of organic solvents in 0.1% trifluoroacetic acid. The organic solvents: (1) 80% Acetonitrile, (2) 50% Acetonitrile, 30% Isopropanol, (3) 50% Acetonitrile, 30% Butanol.

**Supplementary Figure 19**

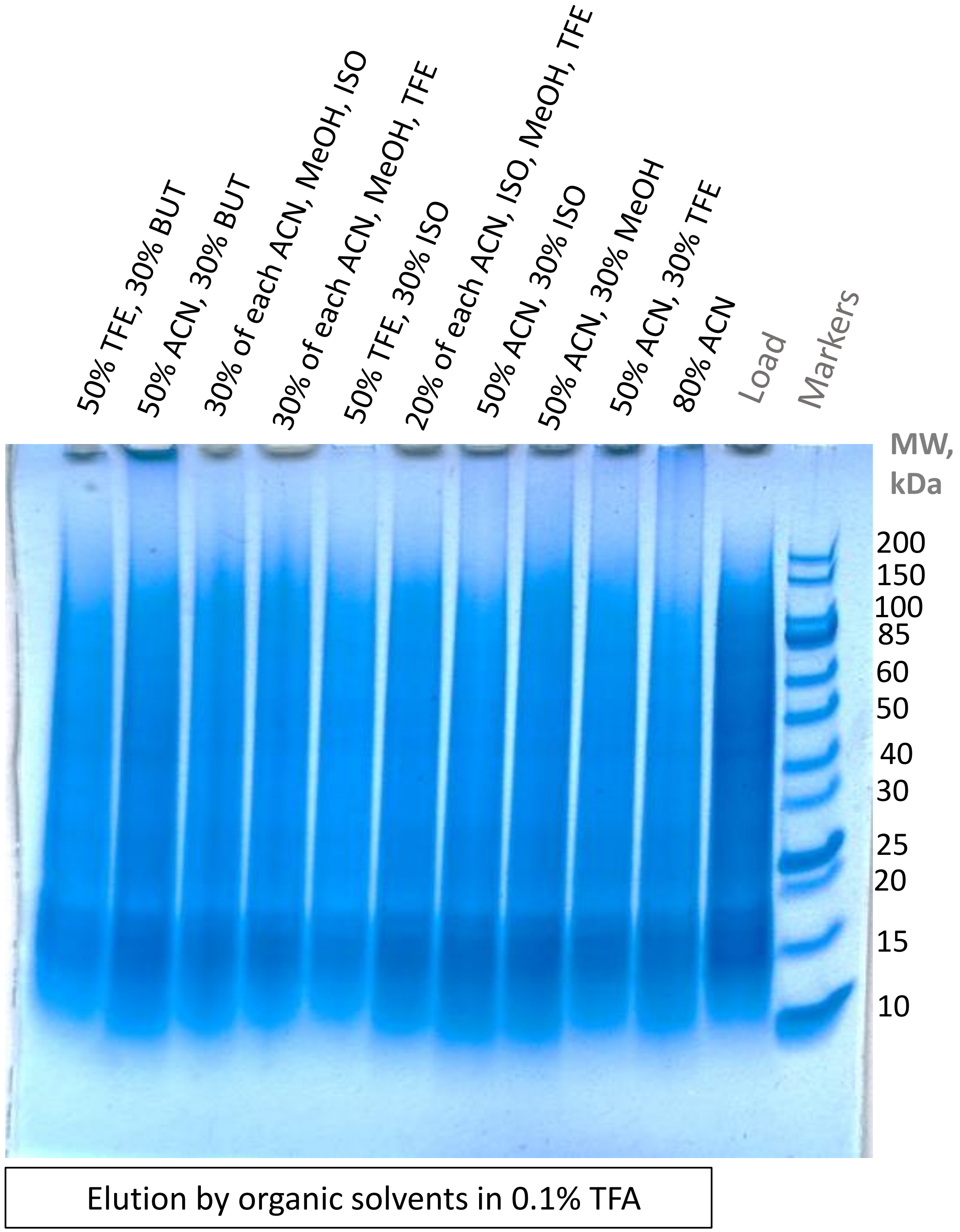


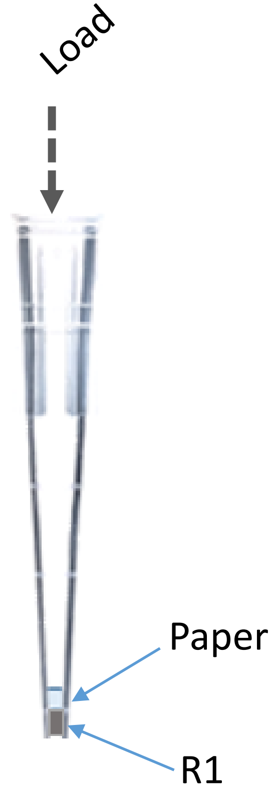


**Protein elution from UMA-R tips using different solvent combinations. COS-7 cell SDS protein lysate was used. Visualization by gel electrophoresis.**

COS-7 cell protein lysate in 4% SDS was loaded into UMA-R tips (GB003 paper part, POROS R1 bottom part) in 6 volumes of the AMA solution (45% acetonitrile, 45% methanol, 100 mM Ammonium Acetate (AA)). After the post-load AMA wash, followed by the wash with 100 mM AA, the paper-captured proteins were eluted onto the R1 reverse phase media after incubation for 2 min at 70 °C with 20% Ethanolamine. Following the wash with 3% acetonitrile, the proteins were eluted from the R1 media with different combinations of organic solvents in 0.1% trifluoroacetic acid (TFA). The organic solvents: ACN – acetonitrile, MeOH – methanol, TFE – trifluoroethanol, ISO – isopropanol, BUT – butanol. The eluates were dried down and reconstituted with 1X Laemmli buffer. The samples were run on a 4-12% Bis-Tris protein gel. The gel was stained with Coomassie.

**Supplementary Figure 20**

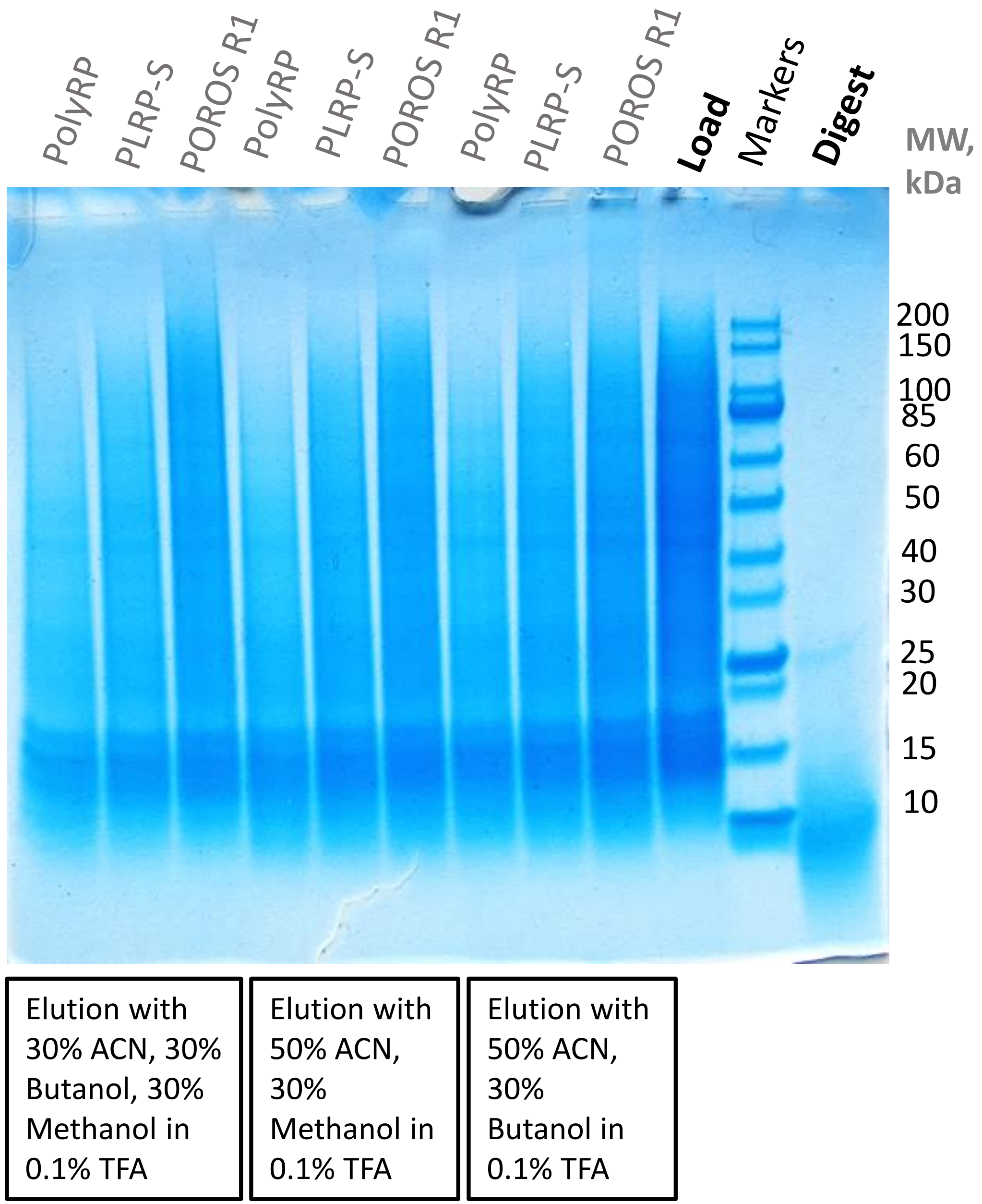


**Testing different RP media (POROS R1, PLRP-S and PolyRP) for protein capture and recovery in UMA-R tips. Protein capture, digest and post-digest peptide recovery using the UMA-R tip with POROS R2 material. COS-7 cell SDS protein lysate was used. Visualization by gel electrophoresis.**

UMA-R tips were constructed using paper and either POROS R1, PLRP-S or PolyRP reversed phase (RP) media. COS-7 cell lysate in 4% SDS was loaded into the tips in 6 volumes of the AMA solution. After the post-load AMA wash, followed by the wash with 100 mM AA, the paper-captured proteins were eluted onto the RP after incubation for 2 min at 70 °C with 20% Ethanolamine. Following the wash with 3% acetonitrile, the proteins were eluted from the RP part with different combinations of organic solvents in 0.1% TFA. The organic solvents: acetonitrile (ACN), methanol and butanol. In addition, the COS-7 cell lysate was loaded onto the UMA-R tip (constructed with paper and POROS R2 media) in excess of the AMA solution, as above, followed by the AMA wash and the wash with 100 mM AA . The digestion for 20 min at 57 °C was performed using trypsin in 70 mM Ammonium Bicarbonate. The R2-captured peptide products were eluted with 60% ACN in 0.2% FA. The eluates were dried down and reconstituted with 1X Laemmli buffer. The samples were run on a 4-12% Bis-Tris protein gel. The gel was stained with Coomassie.

**Supplementary Figure 21**
