## Supplementary Methods for "Unspecific Molecular Adsorption (UMA) sample preparation method for bottom-up and whole protein analysis. The foundation"

### UMA units

UMA mini units were constructed by introducing 5-mm-diameter filter paper discs into mini spin columns. The filter paper used throughout the study was GB003 blotting paper (Whatman). For initial testing, the following filter papers were also used: generic paper napkin, Whatman's filter papers – grade 4, grade 542, and grade 598, as well as Ahlstrom's 6170 filter paper. UMA paper tips used throughout the study were constructed by inserting 2-mm-diameter plugs of the GB003 blotting paper into 200-µl pipette tips. Also, for initial testing, other UMA tip types were constructed by inserting ~2-mm-diameter glass wool (20383, Supelco), generic cotton wool and nylon mesh (mesh 500) plugs into 200-µl pipette tips. UMA pipetting (UMA-P) tips were constructed by inserting ~2-mm-diameter stainless steel mesh material plugs (woven wire, mesh 500, Inoxia Ltd.) into 200-µl pipette tips. For initial testing, UMA pipetting tips were also constructed by inserting 2-mm-diameter generic polyester felt or nickel foam plugs into 200-µl pipette tips. UMA-R tips were constructed by loading wide pore polymeric PS-DVB media - POROS R1, pore size 4000 Å or POROS R2, pore size 2000 Å (Thermo), PLRP-S, pore size 4000 Å (Agilent) or PolyRP, pore size 1000 Å (Sepax) into fritted 200-µl pipette tips and placing either the GB003 filter paper or cotton wool plugs on the top. UMA-$C_{18}$ tips were constructed by loading Bond Elut $C_{18}$ media (Agilent) into fritted 200-µl pipette tips and placing either the GB003 filter paper or cotton wool plugs on the top.

### UMA units with embedded enzymes

UMA tips with embedded enzymes were constructed by inserting two 2-mm-diameter plugs of the GB003 blotting paper (Whatman) or, initially, ~2-mm-diameter generic cotton wool balls into 200-µl pipette tips. Also, to test enzyme binding to non-cellulose materials, 200-µl pipette tips with QM-A silica filter (Whatman) and generic polyester felt 2-mm plugs were made. The UMA tips were washed with deionized water. A solution of an enzyme (~ 2-2.5 µg) dissolved in deionized water at the concentration of ~ 0.01 µg/µl was loaded into the UMA tips from above and slowly pushed down through using a syringe with an appropriate tip adapter. The UMA tips were washed consecutively with deionized water and ethanol (for dehydration). The UMA tips were left to dry out at room temperature (RT) for 24 hours before use or storage. UMA mini units with embedded enzymes were constructed by introducing 5-mm-diameter GB003 filter paper discs with embedded enzymes into mini spin columns. To embed enzymes into a piece of filter paper the following procedure was followed – to a square piece of the GB003 filter paper (10 X 10 cm) 150 µg of an enzyme dissolved in deionized water at the concentration of ~ 0.01 µg/µl was added and incubation for 15 min with mild shaking at RT was performed. The piece of filter paper was flipped over, the enzyme solution was disposed of and a fresh solution of 150 µg of the enzyme dissolved in deionized water at the concentration of ~ 0.01 µg/µl was added followed by incubation for 15 min with mild shaking. After that, the enzyme solution was disposed of, the filter paper was washed consecutively with deionized water and ethanol. The filter paper was then left to dry out at RT for 24 hours prior to use or storage. The following enzymes were used - elastase (V189A, Promega), trypsin (T0303, Sigma) and chymotrypsin (9004-07-3, Merck).

### Cell lysates

HT-29, PC3, COS-7, 293T or MGH-U3 frozen cell pellets (~4,000,000 cells per pellet) were lysed by probe sonication in 500 µl of either 4% SDS, 50 mM Tris-HCl, pH 7.6 or 4% SDS, 50 mM $NaHCO_3$ solutions. The extracts were clarified by centrifugation at 11,000xg for 5 min. The clarified extracts were further filtered through Ultrafree-MC (Amicon) 0.45-µm centrifugal filtration units. Protein concentrations were measured using Pierce BCA Protein Assay Kit (Thermo).

### Dyeing with Remazol Blue dye

To cellular protein lysates in 4% SDS, 50 mM $NaHCO_3$ or to proteins solubilized in the same solution, Remazol Brilliant Blue dye (R8001, Sigma) was added at the dye-to-protein weight ratio of 1:8. This was followed by incubation at 55 °C for 20 hours. The dyed protein solutions were stored at RT.

### UMA processing

For direct loading, to cellular protein lysates or protein solutions, 6 volumes of the AMA loading solution (45% methanol, 45% acetonitrile in 100 mM ammonium acetate) were added. The mixtures were loaded into the UMA mini units or UMA tips either by centrifugation at 2,000xg or by applying positive pressure with a syringe attached to an appropriate adaptor. The captured proteins were washed with the AMA solution and, optionally, with the 70 mM AmBic solution. After the capture and clean-up, an enzyme (e.g. porcine trypsin, T0303, Sigma) dissolved in the 70 mM AmBic solution was applied. The units were incubated at elevated temperatures (53 °C-57 °C) with the incubation time range up to 45 min. The resultant peptide products were eluted for further analysis. For direct loading by pipetting, to cellular protein lysates or protein solutions, from 5 to 6 volumes of the AMA loading solution were added. The mixtures were pipetted through the UMA-P pipetting tips up to 30 times. The captured proteins were washed with the AMA solution. Digestion was performed, similarly as described above, by an applied enzyme. The resultant peptide products were eluted for further analysis.

### UMA processing with embedded enzymes

To cellular protein lysates or protein solutions, 6 volumes of the AMA loading solution (45% methanol, 45% acetonitrile in 100 mM ammonium acetate) were added. The mixtures were loaded into UMA mini units or UMA tips containing embedded enzymes either by centrifugation at 2,000xg or by applying positive pressure with a syringe equipped with an appropriate adaptor. The captured proteins were washed with the AMA solution. Afterwards, the 70 mM AmBic solution was added. The units were incubated at elevated temperatures (53 °C-57 °C) with incubation times from 5 min to 45 min. The resultant peptide products were eluted for further analysis.

### Serum processing

A sample of normal human serum was obtained from the Leeds Multidisciplinary Research Tissue Bank. For fractionation on UMA-P (stainless steel mesh) tips, 1 µl of the serum was diluted in 100 µl of the 250 mM TEAB, 5 mM TCEP, 10 mM IAA buffer solution and incubated at RT for 10 min. The protein solution was then pipetted up and down using an UMA-P (stainless steel mesh) tip thirty times, the tip was removed from the protein solution and washed with 250 mM TEAB (fraction 1 was retained on the tip). After that, 2 volumes of ACN were added to the protein solution and it was pipetted up and down thirty times using a new UMA-P tip, the tip was removed from the protein solution and washed with 70% ACN, 80 mM TEAB (fraction 2 was retained on the tip). Then, another volume of ACN was added and the solution was pipetted again thirty times using a new UMA-P tip, the tip was removed from the protein solution and washed with 80% ACN, 60 mM TEAB (fraction 3 was retained on the tip). The bound proteins were either eluted with 1X Laemmli buffer or digested for 45 min at 53$^{o}$C with applied 2 µg of trypsin in 70 mM AmBic, eluted in 70 mM AmBic, dried down and reconstituted with 0.2% FA for further LC-MS/MS analysis. For unfractionated serum processing, 1 µl of the serum was solubilized in 25 µl of the 4% SDS, 50 mM TEAB solution, consecutively reduced and alkylated with 10 mM DTT and 25 mM IAA. The proteins were loaded into an UMA paper mini unit with 6 volumes of the AMA (45% methanol, 45% acetonitrile in 100 mM ammonium acetate) solution. The captured proteins were washed with the AMA solution. The bound proteins were digested for 45 min at 53$^{o}$C with applied 2 µg of trypsin in 70 mM AmBic, eluted in 70 mM AmBic, dried down and reconstituted with 0.2% FA for further LC-MS/MS analysis.

### Immunoprecipitation

Immunoprecipitation using polyclonal rabbit anti-FOXM1 antibody (13147-1-AP, Proteintech) stored at 4 °C for 10 years and control IgG from rabbit serum (I5006, Sigma) was performed. Six frozen HeLa cell pellets (~4 million cells each) were extracted with 0.5 ml per pellet of the Radio-Immunoprecipitation Assay (RIPA) buffer containing protease inhibitors (Complete Mini, EDTA-free, Roche). The extracts were cleared by centrifugation at 11,000xg for 5 min and then combined. The combined extracts volume was adjusted to 3 ml with the RIPA buffer. Immunoprecipitation was performed in triplicates for the two tested antibodies. A 2-µg antibody aliquot was added to 0.5 ml of the HeLa extract. After rotation at RT for 30 min, 30 µl of Protein G magnetic beads (Dynabeads, Thermo) was added followed by another rotation for 30 min. The extract was removed and the beads were washed with the RIPA buffer. Captured proteins were eluted by incubating the beads with 30 µl of 4% SDS, 50 mM TRIS-HCl, pH 7.6 buffer containing 20 mM DTT at 90$^{o}$C for 5 min. Alkylation was performed by adding IAA to the final concentration of 60 mM and incubation for 30 min at RT in the dark. The samples were processed using tryptic digestion on UMA paper tips.

### Protein acidic cleavage with TCEP reduction

Bovine insulin (I6634, Sigma), porcine trypsin (T0303, Sigma), bovine carbonic anhydrase (C3934, Sigma) or human serum albumin (A3782, Sigma) were separately solubilized in the 4% SDS, 50 mM TEAB buffer. 20 µg of solubilized protein was loaded in 6 volumes of the AMA buffer into an UMA-$C_{18}$ tip. The captured proteins were washed with the AMA solution and deionized water. The acidic solution of 10% HCl, 50 mM TCEP was then applied, the tips were capped on the top and placed at 85 °C for 30 min. The acidic solution was removed, the tips were washed with deionized water and the peptides were eluted with 60% ACN, 0.2% FA. The eluted peptides were dried down and reconstituted in 1% ACN, 0.2% FA for further LC-MS/MS analysis.

### Protein acidic cleavage without TCEP reduction

Human serum albumin (A3782, Sigma) was solubilized in the 4% SDS, 50 mM TEAB buffer and consecutively reduced and alkylated with 20 mM DTT and 50 mM IAA. Sample processing was done similarly to the described above except that the applied acidic cleavage solution contained only 10% HCl.

### Protein digest with the embedded elastase

Bovine insulin (I6634, Sigma), porcine trypsin (T0303, Sigma), bovine carbonic anhydrase (C3934, Sigma) or human serum albumin (A3782, Sigma) were separately solubilized in 4% SDS, 50 mM TEAB buffer and consecutively reduced and alkylated with 20 mM DTT and 50 mM IAA. 20 µg of solubilized protein was loaded in 6 volumes of the AMA solution into an UMA paper tip with embedded elastase. The captured proteins were washed with the AMA solution. 70 mM AmBic was then applied and the tips were placed at 57 °C for 30 min. The peptides were eluted in 70 mM AmBic, followed by 30% ACN, 0.2% FA. The eluted peptides were dried down and reconstituted in 1% ACN, 0.2% FA for further LC-MS/MS analysis.

### MS analysis of tryptic digests on Orbitrap Exploris 240

Concentration of the digest products was directly measured on a NanoDrop spectrophotometer (Thermo) by utilizing the absorbance at 280 nm. ~3 µg of peptides were loaded and separated by reversed-phase capillary chromatography using an EASY-nLC (Thermo) unit connected to a 15-cm capillary emitter column (75 µm inner diameter, 3 µm Reprosil-Pur 120 $C_{18}$ media, Dr. Maisch). The LC system was hyphenated with an Orbitrap Exploris 240 mass spectrometer (Thermo). The acquisition time per run was 45 min, the main part of the gradient was 4% - 25% acetonitrile in 0.1% formic acid. The MS scan resolution was set to 60,000 and the m/z scan range was 300 – 1350 amu. Up to 20 most intense multiply charged ions per scan were fragmented in the MS2 mode, the MS2 resolution was set to 15,000. Either single or duplicate LC-MS/MS sample runs were performed. Data processing against a Uniprot human protein sequence database (August, 2024) and label free quantitation were done with MaxQuant 2.4.13.0 software package (www.maxquant.org). Carbamidomethylation of cysteine was chosen as a fixed modification, and protein N-terminal acetylation, methionine oxidation, and asparagine and glutamine deamidation set as variable modifications. At least one unique peptide for valid protein identification was chosen. Besides that, the default software settings were used.

### MS analysis of protein cleavage products on LTQ Orbitrap Velos

To analyze the elastase protein digest, protein acidic cleavage with incorporated TCEP reduction or protein acidic cleavage, the peptide loading and separation were done similarly to the described above. The acquisition time per LC-MS/MS run was 35 min, the main part of the gradient was 4% - 25% acetonitrile in 0.1% formic acid. The EASY-nLC system was hyphenated with an LTQ Orbitrap Velos (Thermo) mass spectrometer. MS scans (330 – 1500 amu range) were acquired in the orbitrap with the resolution set to 60,000. Up to 20 most intense multiply charged ions per scan were fragmented in the linear ion trap. Data were searched against a Uniprot protein sequence database with Mascot search engine (Matrix Science) using the 'no enzyme' unspecific cleavage option. Carbamidomethylation of cysteine was chosen as a fixed modification for the samples where the elastase digest was performed (the samples had been reduced and alkylated with iodoacetamide prior to the elastase digest), and protein N-terminal acetylation, methionine oxidation, and asparagine and glutamine deamidation were set as variable modifications for the elastase digest and reductive acidic cleavage. For the sequence database search of proteins reduced and alkylated with iodoacetamide prior to their acidic cleavage (without the incorporated TCEP reduction) the following variable modifications were chosen - protein N-terminal acetylation, methionine oxidation, asparagine and glutamine deamidation, and cysteine carboxymethylation. The significance threshold was set to $p<0.05$ and the ion score cut-off was set to 10.

### LC-MS profiling of protein mixture on LTQ Orbitrap Velos

Bovine insulin (I6634, Sigma), porcine trypsin (T0303, Sigma), bovine carbonic anhydrase (C3934, Sigma) and human serum albumin (A3782, Sigma) were solubilized in 4% SDS, 50 mM TEAB buffer with the weight-to-weight ratio 1:4:5:10, respectively. 20 µg of this protein mixture was processed on an UMA paper tip via the loading in 6 volumes of the AMA solution. After the wash with the AMA solution, the proteins were eluted with 35% TFE, 1% FA. ~2 µg of the eluted protein mixture was loaded and separated by chromatography using an EASY-nLC LC (Thermo) unit connected to a 15-cm capillary emitter column (150 µm inner diameter, 5 µm ReproSil-XR 300 $C_4$ media, Dr. Maisch). The LC system was hyphenated with an LTQ Orbitrap Velos mass spectrometer. MS scans (scan range 500-2000 amu) were acquired in the orbitrap with the resolution set to 100,000. The acquisition time per run was 35 min, the main part of the gradient was 5% - 80% of the 50% acetonitrile, 30% isopropanol, 0.1% trifluoroacetic acid elution solution.

### Antibody profiling by LC-MS on BioAccord

A commercial monoclonal antibody (A2228, Sigma, MW ~150 kDa) was either used as is or had SDS added to it to the final 4% concentration. The antibody samples were loaded onto the UMA paper tips in 6 volumes of the AMA solution, washed with the AMA solution and eluted after 2-min incubation in the 0.1% TFA elution solution at RT. The mass spectrometric profiling of the antibody eluates was done using reversed phase chromatographic separation on the MAbPac phenyl column (catalog number 088647, 2.1x100 mm, Thermo) and BioAccord LC-MS TOF System (Waters). ~1 µg of the eluted protein was loaded onto the column. The total acquisition time per run was 14 min, the 7-minute gradient was 20% - 80% of the acetonitrile in 0.1% formic acid. The TOF MS scans (scan range 400-7000 amu) were acquired with default resolution of 20,000.

### Protein profiling by ESI-MS

Carbonic anhydrase (C3934, Sigma), alcohol dehydrogenase yeast (A7011, Sigma) or myelin basic protein (M1891, Sigma) were solubilized separately in the 4% SDS, 50 mM TEAB buffer. Additionally, alcohol dehydrogenase was reduced with DTT and alkylated with iodoacetamide. 20 µg of solubilized protein was cleaned up using the AMA loading and either UMA tip with a cotton wool plug, UMA tip with a paper plug or UMA-P pipetting tip with a felt plug. After the wash with the AMA solution, the proteins were eluted either with 35% TFE, 1% FA or 70% TFE, 1% FA. The spectra were acquired on a Synapt G1 hybrid quadrupole-time-of-flight instrument (Waters) utilizing coated borosilicate capillary emitters, with the MS scans within the 500-4000 amu range and default resolution of 10,000.

### Protein processing on UMA-R tips

Dyed or undyed protein mixtures were loaded onto UMA-R tips (the tip's upper part – paper or cotton wool, the tip's bottom part – wide pore PS-DVB beads) in 6 volumes of the AMA solution. The UMA-R tips were washed with the AMA solution followed, optionally, by the wash with 100 mM ammonium acetate. The following protein solubilization solutions were loaded into the UMA-R tips - 50% FA or 70% TFE, 100 mM TEAB or 20% Ethanolamine followed by incubation for 2 min at RT or, optionally, at 70 °C (this optional incubation at elevated temperatures, e.g. 70 °C, may help in solubilization of the captured proteins). The solubilized proteins were then forwarded onto the PS-DVB bottom part of the UMA-R tips, washed with 3% ACN and eluted by concentrated acidic acetonitrile or acetonitrile-alcohol mixtures (e.g. 50% ACN, 30% ISO, 0.1 % TFA).

### Tips with ion-exchange media

SCX tips were constructed by loading Bio-Pro S75 media (YMC) into fritted 200-µl pipette tips and placing the GB003 filter paper (Whatman) plugs on the top. SG81 tips were constructed by inserting the SG81 silica paper (Whatman) plugs into 200-µl pipette tips and placing the GB003 filter paper plugs on the top.  SG81 mini units were constructed by introducing the 5-mm-diameter SG81 silica paper discs into mini spin columns. Quartz tips were constructed by inserting the QM-A quartz (Whatman) plugs into 200-µl pipette tips and placing the GB003 filter paper plugs on the top. The SCX, SG81 and quartz units were used to capture/process the proteins solubilized with the acidic TFE solution (70% TFE, 1% FA).
